# Vhl deletion in *Dmp1*-expressing cells alters MEP metabolism and promotes stress erythropoiesis

**DOI:** 10.1101/2023.07.25.550559

**Authors:** Janna M. Emery, Betsabel Chicana, Hanna Taglinao, Citlaly Ponce, Cristine Donham, Hawa Padmore, Aimy Sebastian, Scott L. Trasti, Jennifer O. Manilay

**Author notes:** Joint co-first authors (Janna M. Emery, Betsabel Chicana). Correspondence: Jennifer O. Manilay, PhD; University of California Merced, Merced, CA 95343.

## Abstract

In recent years, general hypoxia-inducible factor (HIF)-prolyl hydroxylase (PHD) enzyme inhibitors have been developed for the treatment of anemia due to renal disease and osteoporosis. However, it remains a challenge to target the HIF signaling pathway without dysregulating the skeletal and hematopoietic system. Here, we examined the effects of *Vhl* deletion in bone by performing longitudinal analyses of *Vhl*cKO mice at 3, 6, 10, and 24 weeks of age, where at 10 and 24 weeks of age, high bone mass and splenomegaly are present. Using flow cytometry, we observed increased frequency (%) of CD71^lo^TER119^hi^FSC^lo^ orthochromatophilic erythroblasts and reticulocytes in 10- and 24-week-old *Vhl*cKO bone marrow (BM), which correlated with elevated erythropoietin levels in the BM and increased number of red blood cells in circulation. The absolute numbers of myeloerythroid progenitors (MEPs) in the BM were significantly reduced at 24 weeks. Bulk RNA-Seq of the MEPs showed upregulation of *Epas1* (*Hif1a)* and *Efnb2* (*Hif2a)* in *Vhl*cKO MEPs, consistent with a response to hypoxia, and genes involved in erythrocyte development, actin filament organization, and response to glucose. Additionally, histological analysis of *Vhl*cKO spleens revealed red pulp hyperplasia and the presence of megakaryocytes, both of which are features of extramedullary hematopoiesis (EMH). EMH in the spleen was correlated with the presence of mature stress erythroid progenitors, suggesting that stress erythropoiesis is occurring to compensate for the BM microenvironmental irregularities. Our studies implicate that HIF-driven alterations in skeletal homeostasis can accelerate erythropoiesis.

**Key Points:** • Dysregulation of HIF signaling in Dmp1+ bone cells induces stress erythropoiesis.

• Skeletal homeostasis modulates erythropoiesis.

## INTRODUCTION

Erythroid homeostasis is highly regulated to maintain enough red blood cells (RBC) for tissue oxygenation and avoid their overproduction, which causes viscosity-related complications. RBCs are derived from progenitors that reside in the bone marrow (BM). To produce RBC, megakaryocyte-erythroid progenitors (MEP) differentiate into burst forming unit-erythroid and colony-forming unit-erythroid (CFU-E)^1^. With erythropoietin (EPO) stimulation, CFU-E develops into various erythroid progenitors: proerythroblasts, basophilic erythroblasts, polychromatophilic erythroblasts, and orthochromatophilic erythroblasts^2,3^. These progenitors develop in proximity with macrophages in a specialized BM niche called an erythroblastic island^4^. To develop mature RBCs, terminal maturation involving enucleation to form reticulocytes occurs^5^. Reticulocytes then undergo organelle clearance and membrane remodeling^5,6^.

Although steady state erythropoiesis has the immense capacity to produce new RBC, there are instances when it is unable to maintain erythroid homeostasis. In these times, a short-term, compensatory mechanism termed stress erythropoiesis produces a bolus of new RBC to maintain homeostasis until steady state erythropoiesis can resume^7^. This response is driven by stress erythroid progenitors (SEPs) and is best understood in mice, where it is extramedullary, occurring in a specialized microenvironment in the fetal liver and both the adult spleen and liver^8–10^. While extensive efforts have been made to understand RBC development, many questions remain regarding how the BM and bone microenvironment maintains and regulates myeloerythroid development.

Bone is a highly dynamic tissue that undergoes continuous remodeling to maintain a healthy skeleton, which is important for supporting efficient and lifelong skeletal functions. Dysregulation of coupled signaling pathways or imbalances in bone resorption and formation can lead to abnormal bone remodeling and the development of bone diseases. Osteoporosis is a bone disease characterized by low bone mass and density, causing bone fragility and an increased risk of fractures^11^. Current strategies involving the usage of osteolineage-specific *Cre* drivers to delete of either prolyl hydroxylase (PHD) or von Hippel Lindau (VHL) and stabilize hypoxia inducible factor (HIF) result in increased bone mass, leading to a novel, targeted basis of treating bone loss in individuals with osteoporosis using PHD and VHL inhibitors^12–15^. However, the concerns of modifying inhibitors or activators of the HIF pathway remain, due to potential adverse effects on the immune system^16^.

In this study, we utilized *Dmp1*-Cre; *Vhl* conditional knockout mice (*Vhl*cKO) mice, where *Vhl* is deleted in subsets of mesenchymal stem cells, late osteoblasts, and osteocytes^17^ to investigate how changes in bone homeostasis affects myeloerythroid development. *Vhl*cKO mice present with high bone mass that results in a severely occluded BM cavity with reduced BM cellularity, dysregulated B cell development, and prominent splenomegaly^14,16^. Here, we provide evidence for structural, molecular, and cellular alterations in the *Vhl*cKO BM niche that inadvertently alters the state of erythropoiesis, such as gene expression changes in MEPs and increased EPO levels. We also observed an expansion of the erythroid niche to the spleen that could be functioning to compensate for the BM irregularities. These studies reveal novel mechanisms by which *Vhl* deletion in *Dmp1*-expressing cells can modulate erythropoiesis.

## METHODS

### Experimental Animals

Age-matched male and female mice on the C57BL/6 (B6) background were used. *Dmp1*- Cre (JAX 023047) and *Vhl^fl/fl^* (JAX 012933) mice were crossed to generate conditional *Vhl* knockouts (*Vhl*cKO) in *Dmp1*-expressing cells^18,19^. Genotyping was confirmed by PCR^16^ or Transnetyx, Inc. (Cordova, TN) using real-time PCR and no sex-specific differences in erythropoiesis or other components of our studies were detected. Mice were housed and bred under specific-pathogen free conditions. The University of California, Merced Institutional Animal Care and Use Committee approved all animal work.

### Bone Marrow and Spleen Collection and Processing

Mice were euthanized by CO_2_ asphyxiation followed by cervical dislocation. BM from long bones and splenocytes from spleens were isolated as described^16^. Isolated cells were treated with ACK lysis buffer to remove erythrocytes for myeloid progenitors, lineage, and erythroblast analysis. Cell counts were obtained using a hemocytometer and Trypan Blue staining to exclude dead cells.

### Flow Cytometry

Peripheral blood was collected into Eppendorf tubes with heparin (Sigma-Aldrich, Inc., St. Louis, MO) and hypotonic lysis of RBCs was performed^20^. Cell staining included a pre-incubation step with unconjugated anti-CD16/32 (clone 93) to block Fc receptors, except for myeloid progenitor panel^16^. The antibody cocktails used are listed in **Supplemental Table 1**. For viability staining, DAPI (Sigma-Aldrich, 1 μg/ml) or PI (Sigma-Aldrich, 1 μg/ml) was used. Single color stains were used for setting compensations and gates were determined with fluorescent-minus one controls, isotype-matched antibody controls, or historical controls. Data were acquired with BD LSR II (Becton-Dickinson) or ZE5 (BioRad, Hercules, CA) flow cytometers. The data was analyzed using FlowJo Software version 10.7.1.

### Bulk RNA-Sequencing

MEPs (live, negative for F4/80, CD3, CD4, CD5, CD8, CD19, NK1.1, Ter119 and Gr1; CD45+ cKIT+ Sca1-CD34-CD16/32-) from pooled BM of control and *Vhl*cKO mice were isolated by flow cytometric sorting on the FACS Aria3 (Becton-Dickinson) and stored in RLT lysis buffer (Qiagen, Redwood City, CA) supplemented with β-mercaptoethanol at -80C until RNA isolation. RNA from sorted MEPs was isolated using RNEasyMinElute columns following the manufacturer’s instructions. The isolated RNA from each sample was flash frozen and sent to UC Irvine Genomics Research and Technology Hub on dry ice to obtain a library of transcripts. Library construction was performed with the SMARTer® Stranded Total RNA-Seq Kit v3 and sequenced on the i5 NovaSeq 6000. Sequencing data quality was checked using fastQC^21^. Subsequently UMIs, adapters and linkers were trimmed from Read 2 using Trimmomatic^22^. Reads were then mapped to mouse genome (mm10) using STAR^23^ and read counts per gene were determined using “featureCounts” from Rsubread package^24^. Differentially expressed genes (DEGs) were identified using limma after voom normalization^25^. Genes with a P value <0.05 and fold change > 2 were considered as significantly differentially expressed. Result visualizations were generated using custom scripts conducted in R version 4.3.0 (2023-04-21)^26^. Functional gene ontology was analyzed from DEGs in the ToppGene Suite for gene list enrichment analysis with a false discovery rate set to p < 0.05^27^. Sequence files were submitted to the Gene Expression Omnibus (GEO) at the National Center for Biotechnology Information (NCBI) under accession number GSE237723.

### PCR

Whole BM cells were isolated from long bones as described above. Cells were pelleted and resuspended in RNeasy RLT Lysis Buffer (Qiagen) with 1% 2-mercaptoethanol. Total RNA was purified using the Qiagen RNeasy Micro Kit (Qiagen) according to manufacturer’s protocol. RNA concentration and purity was analyzed using the Nanodrop One Spectrophotometer (Thermo Fisher Scientific). For RT-qPCR, the same amount of RNA from each sample was mixed with qScript XLT One-Step, RT-qPCR ToughMix (Quantabio) together with specific TaqMan expression primers. Real-time qPCR was run on the Applied Biosystems thermocycler using QuantStudio3 software (ThermoFisher) at the following specifications: 1 cycle at 50°C for 10 minutes for cDNA synthesis, 1 cycle at 95°C for 1 minute for initial denaturation and then 40 cycles of amplification at 95°C for 5 seconds then 60°C for 45 seconds. The following TaqMan gene expression assays (Thermo Fisher Scientific) were used: housekeeping *Actb*-VIC (Mm02619580_g1), target genes *Epo*-FAM (Mm01202755_m1) and *EpoR*-FAM (Mm00833882_m1).

For reverse transcriptase-PCR, a 5 mg piece of kidney tissue was homogenized in 350 *μ*L of RNeasy RLT Lysis buffer (Qiagen) containing 2-mercaptoethanol and centrifuged at maximum speed for 3 minutes at room temperature. Total RNA was purified using the Qiagen RNeasy Micro Kit (Qiagen). Total RNA concentration and purity was analyzed using a NanoPhotometer (Implen, Westlake Village, CA). Complementary DNA was reverse transcribed using the iScript cDNA Synthesis Kit (BioRad). RT-PCR reactions were run on the SimpliAmp thermocycler (Applied Biosystems, Foster City, CA) using an equivalent to 100 ng of RNA and specific primers for GAPDH2: forward 5’-TCACCACCATGGAGAAGGC-3’, reverse: 5’-GCTAAGCAGTTGGTGGTGCA-3’; and VHL 2-3: forward 5’-GCCTATTTTTGCCAACATCACA-3’, reverse: 5’-CAAGGCTCCTCTTCCAGGTG -3’.

### Quantification of Cytokines

BM fluid and PB was collected and processed, and cytokine measurements were performed using a customized bead-based multiplex (13-LEGENDplex assay) from Biolegend, Inc. as described^16^.

### Complete Blood Count and Peripheral Blood Smears

Tail bleeds were performed as described^16^. For complete blood count (CBC) and PB smears, peripheral blood was directly collected into BD Microcontainer tubes with K2E (K_2_EDTA; Fisher Scientific, Pittsburgh, PA) and 1.5 mL Eppendorf tubes with EDTA, respectively. A CBC was performed using a HEMAVET HV950 automated veterinary hematology counter (Drew Scientific, Inc., Miami Lakes, FL). MULTI-TROL mouse control blood (Drew Scientific, Inc., Miami Lakes, FL) was run prior to mouse blood samples to calibrate the HEMAVET system and assess sample quality control. The quality control program was run daily.

PB smears were prepared within 1 hour of blood collection to minimize artifacts. A glass slide was positioned so that the frosted end was on the left and one drop of blood was placed in the center nearest to the frosted end. Using a second glass slide positioned at a 45° angle, the drop of blood was gently smeared towards the end of the slide and left to air dry for 1 minute. The air-dried blood smear was fixed in methanol, air dried, stained with Wright-Giemsa stain (Volu-Sol, Salt Lake City, UT), and rinsed, then staining was repeated. Stained blood smears were left to air dry and cover slips were mounted using Permount (Fisher Scientific) before microscopic analysis. Slides were shipped overnight to Texas Tech University Health Sciences Center and photomicrographs were taken using a Nikon Eclipse Ni microscope.

### Body Weight and Glucose Measurements

For PB and BM fluid glucose measurements, mice were transferred into a sterile cage with only water for a 12-hour fast. After the 12-hour-fast, body weights (g) were recorded, and PB glucose levels were measured from 1 drop of blood using a glucometer (OneTouch Ultra2 LifeScan).

### Histology

Dissected spleens, livers, and sterna were fixed in 10% neutral buffered formalin (Thermo-Fisher Scientific) for 24 hours. After fixation, all samples were shipped overnight to Texas Tech University Health Sciences Center. Liver and spleen samples were trimmed while sternum samples were decalcified in Cal-Rite solution (Thermo-Fisher Scientific), then placed into histology cassettes. Tissues were processed routinely for histopathology and slides were stained with Hematoxylin and Eosin. Slides were examined and photomicrographs were taken using a Nikon Eclipse Ni microscope.

### Statistical Analysis

A G*Power statistical power analysis (α=0.05 and power of 0.95)^28^ based on myeloid progenitor data and total BM cellularity determined that a minimum of n=8 mice per group was needed for our studies. The total sample size for each experiment was ≥8 performed in three independent experiments. For qPCR of sorted myeloid progenitors, mice were pooled n=2 and ran in duplicates and for whole BM PCR we ran n=4 mice samples in singlets. Age-matched control and *Vhl*cKO mice of both sexes were used. Comparisons between groups were performed using a two-tailed Student’s t-test to test differences between mean and median values with Graph-Pad Prism and were considered significant if p<0.05.

## RESULTS

### Vhl deletion in Dmp1-expressing cells leads to increased myelopoiesis in the bone marrow

Long bones^16^ and sternum from control and *Vhl*cKO mice were collected. *Vhl*cKO bones displayed abnormally high bone mass and occlusion of the BM cavity starting at 6 weeks of age (**Supplemental Figure 1**), which became severe by 24 weeks of age (**Figure 1A**). We previously reported increased frequencies of BM monocytes and granulocytes, suggesting skewing of hematopoietic differentiation towards the myeloid lineages in *Vhl*cKO mice^16^. To elucidate any changes to the myeloid progenitor compartments in the BM of *Vhl*cKO mice, we performed a longitudinal quantification of common myeloid progenitors (CMP: CD45^+^Lin^-^cKIT^+^Sca1^-^ CD16/32^+^CD34^+^), granulocyte/monocyte progenitors (GMP: CD45^+^Lin^-^cKIT^+^Sca1^-^CD16/32^-^ CD34^+^) and megakaryocyte/erythroid progenitors (MEP: Lin^-^CD45^+^cKIT^+^Sca1^-^CD16/32^-^CD34^-^) were analyzed using flow cytometry^29^ (**Figure 1B**). There were no differences at 3 weeks of age (**Figure 1C**). However, starting at 6 weeks of age, *Vhl*cKO mice exhibited an increase in CMP and MEP frequency (**Figure 1C**). By 24 weeks of age, in addition to increased CMP and MEP frequency, these mice also exhibited increased GMP frequency (**Figure 1C**). Absolute numbers of CMPs, MEPs and GMPs were comparable at 3 weeks of age. At 6 weeks, *Vhl*cKO GMPs were reduced in number, but CMPs and MEPs were similar to controls; GMPs then normalized to control levels by 10 weeks of age. By 24 weeks of age, absolute numbers of CMPs, MEPs and GMPs significantly decreased in the *Vhl*cKO BM (**Figure 1C**).

**Figure 1.**
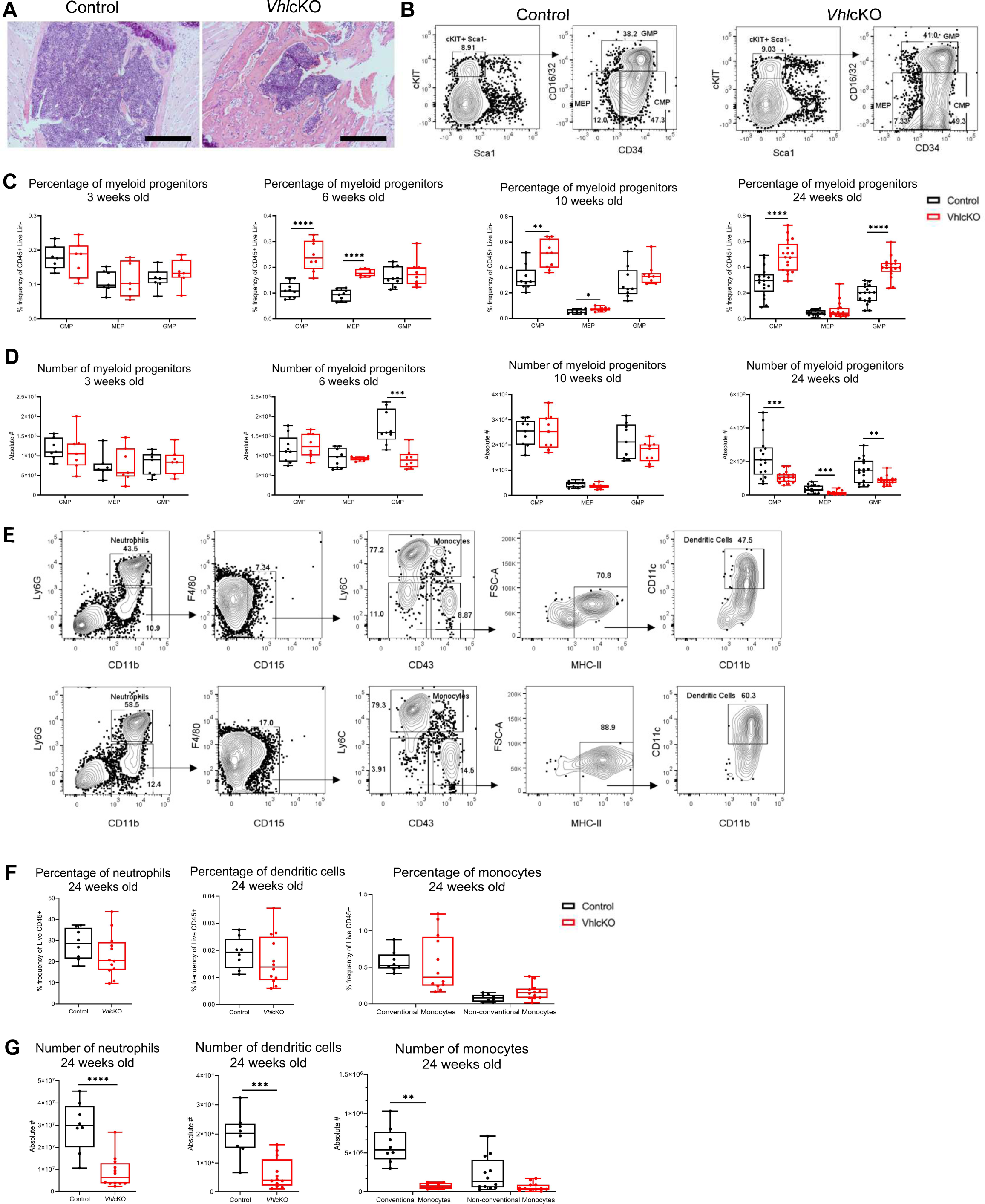
Altered myeloid populations in the bone marrow of *Vhl*cKO mice. **A)** Photomicrographs of hematoxylin and eosin-stained control and *Vhl*cKO sternum, where bone is light pink colored and bone marrow is dark purple colored; scale bar: 100 µm; **B)** Flow cytometry gating strategy for myeloid progenitors; **C)** Percentage and **D)** Number of myeloid progenitors CMPs, MEPs, and GMPs out of Live, CD45+ from bone marrow, by mouse age; **E)** flow cytometry gating strategy of BM myeloid lineage populations; **F)** Percentage and **G)** Number of neutrophils, monocytes, dendritic cells in *Vhl*cKO mice bone marrow. * p < 0.05 ** p < 0.01 *** p < 0.001 **** p < 0.0001, two-tailed Student’s t-test.

We also quantified neutrophils (CD45^+^Ly6G^+^CD11b^+^) classical monocytes (CD45^+^Ly6G^-^ CD11b^+^F4/80^-^CD115^+^Ly6C^hi^) divided into CD43+ or CD43-populations, non-classical monocytes (CD45^+^Ly6G^-^CD11b^+^F4/80^-^CD115^+^Ly6C^-^CD43^+)^, and dendritic cells (CD45^+^Ly6G^-^ CD11b^+^F4/80^-^CD115^+^Ly6C^-^CD43^+^MHC-II^+^CD11c^+^) ^30^ (**Figure 1D**). There were no differences observed in the frequency of neutrophils, monocytes, or dendritic cells in the BM of 24-week-old *Vhl*cKO mice (**Figure 1E, Supplemental Figure 2A-C**). Yet, we observed a decrease in the number of neutrophils, dendritic cells, and conventional monocytes in the BM of 24-week-old *Vhl*cKO mice (**Figure 1F**), consistent with the total decrease in BM cellularity in *Vhl*cKO mice^16^.

### Vhl deletion in Dmp1-expressing cells dysregulates steady state erythropoiesis

The increased frequency of CMPs and MEPs suggested that the state of erythropoiesis may be altered. To address any defects in erythropoiesis, we analyzed erythroblast development in the BM of *Vhl*cKO mice using flow cytometry^2^. We utilized CD71 and TER119 as cell surface markers and FSC as an additional parameter to classify all TER119^+^ cells into four subsets^31^: proerythroblasts (ProE; CD71^hi^TER119^int^FSC^hi^), basophilic erythroblasts (BasoE; CD71^hi^TER119^hi^FSC^hi^), polychromatophilic erythroblasts (PolyE; CD71^lo^TER119^hi^FSC^lo^), and orthochromatophilic erythroblasts and reticulocytes (OrthoE and Retic; CD71^lo^TER119^hi^FSC^lo^) (**Figure 2A**). The absolute number and frequency of all erythroblast populations were unchanged or comparable between 3-week-old control and *Vhl*cKO mice (**Figure 2B, C**). Starting at 10 weeks of age, the frequency of BasoE and PolyE were reduced. However, the frequencies of OrthoE and Retic were increased starting at 6 weeks of age (**Figure 2B**), suggesting that erythropoiesis may be dysregulated in the BM.

**Figure 2.**
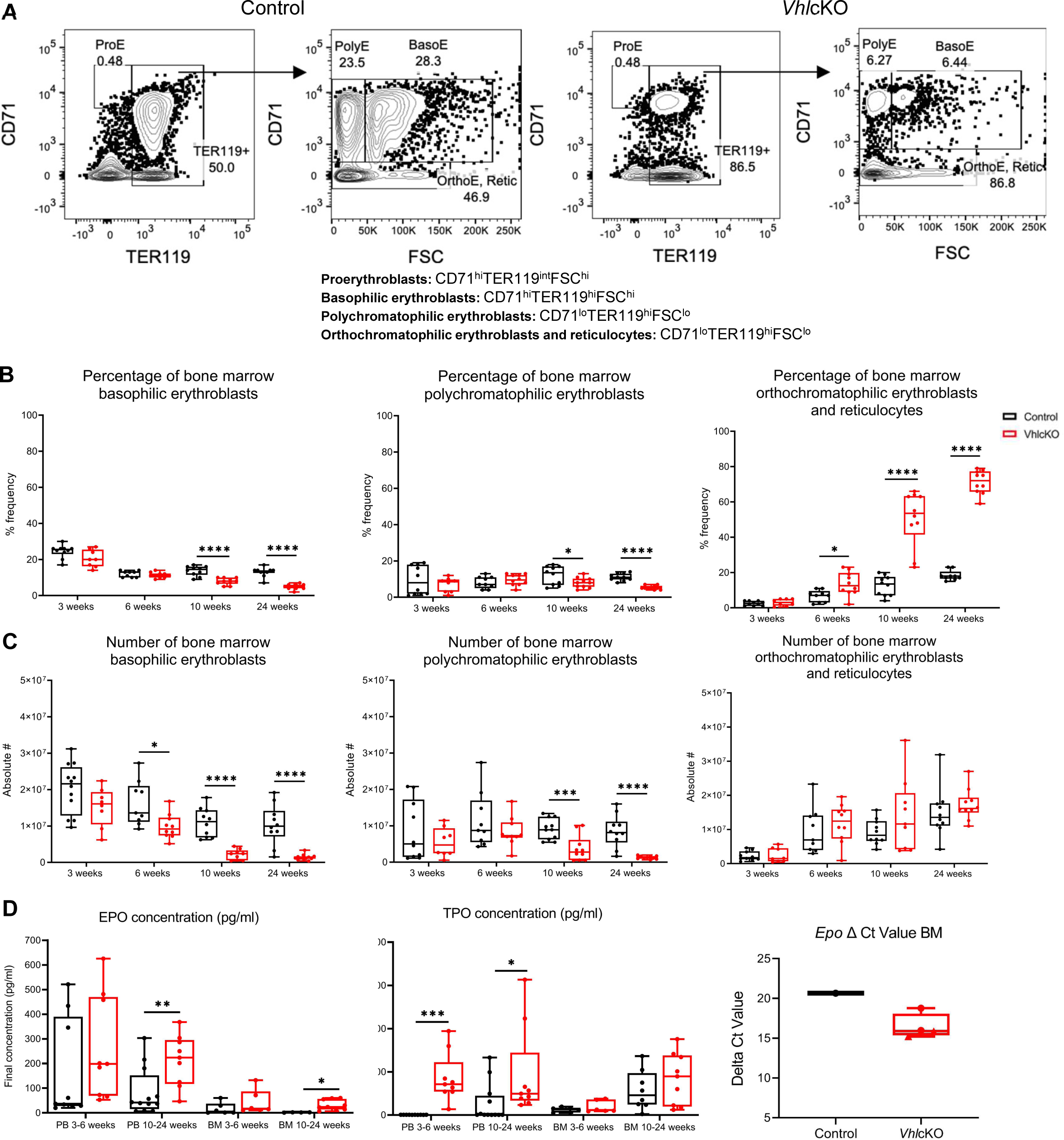
Altered steady state erythropoiesis in the bone marrow of 10- and 24-week-old *Vhl*cKO mice. **A)** Flow cytometry gating of erythroblast populations in the bone marrow of 24-week-old control and *Vhl*cKO mice. ProE, proerythroblasts; BasoE, basophilic erythroblasts; PolyE, polychromatophilic erythroblasts; OrthoE and Retic, orthochromatophilic erythroblasts and reticulocytes; **B)** Percentage and **C)** Number of erythroblast populations; **D)** EPO and TPO protein levels **E)** EPO delta C_t_ value (difference between target gene and housekeeping C_t_ values). * p < 0.05 ** p < 0.01 *** p < 0.001 **** p < 0.0001, two-tailed Student’s t-test.

### Elevated levels of Glut1 and downregulation of EpoR in VhlcKO myeloid progenitors

We next sorted MEPs for bulk RNA sequencing analysis and identified 419 differentially expressed genes (up: 414 genes; down: 5 genes) in *VhlcKO* mice compared to controls (**Figure 3A**, **Supplemental Figure 3, Table 2**). *Epas1* (*Hif1a)* and *Efnb2 (Hif2a)* were upregulated in *Vhl*cKO MEPs, consistent with a response to hypoxia, and levels of *Vhl* expression confirming no off-target deletion (**Figure 3B**). Additionally, we observed an upregulation of genes involved in erythrocyte development (*Epb41l3, Epb41l1, F2rl1*) and actin filament organization (*Myo7b, Shroom2, Cobl, Vil1, Neb*) (**Figure 3B-C**). Moreover, *Slc2a1* (aka *Glut1*) and *Pck1*, key genes involved in response to glucose and metabolic regulation were upregulated and increased *Glut1* expression in MEPs was confirmed by qPCR (**Figure 3D-F**). Knowing that Epo-receptor (EpoR) signaling mediates cell metabolism through an increase in glycolysis^32^, we hypothesized that the increased *Glut1* expression is EpoR mediated. Sequencing results of sorted MEPs revealed a downregulation of *EpoR*, which is consistent with our whole BM qPCR results, but not with our sorted MEP qPCR results, which showed no significant changes (**Figure 3D-F)**. These findings suggest that *Vhl* deletion in *Dmp1*-expressing cells may increase glucose uptake in MEPs and directly affect myeloerythroid development in the BM.

**Figure 3.**
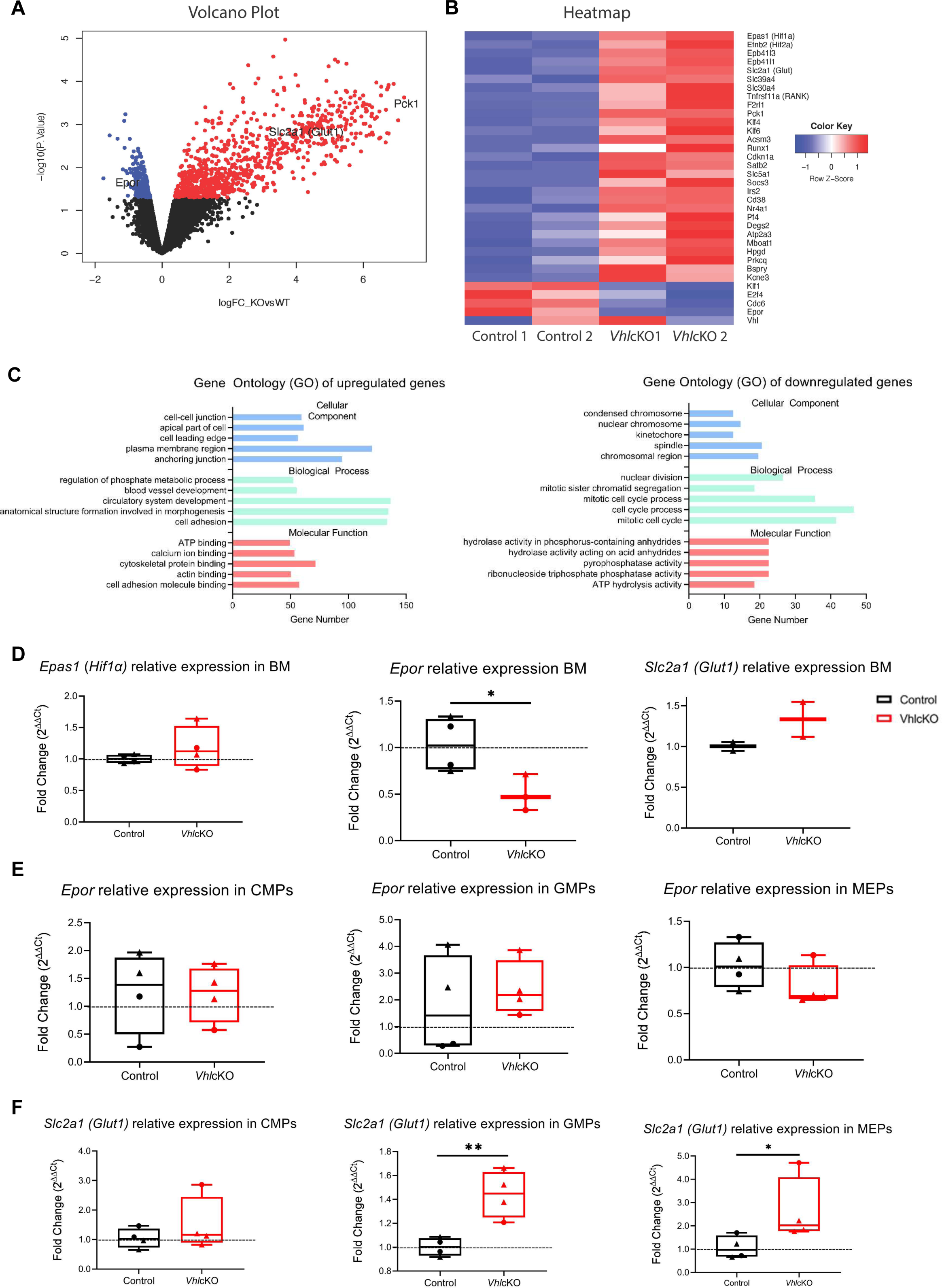
Gene expression analysis of *Vhl*cKO myeloid progenitors. **A)** Volcano plot, fold change (x axis) vs statistical significance (y axis) of upregulated genes (red) and downregulated genes (blue) p<0.05, black represents genes that were not found to be significantly differentiated between *Vhl*cKO and control samples; **B)** Heatmap of 34 significant genes of interest from control and *Vhl*cKO n=2 raw counts of sequenced samples; **C)** Gene Ontology (GO) classification showing significant enrichment of three main gene pathway categories (biological process, cellular component, and molecular function) from bulk RNA sequencing of sorted MEPs from control and *Vhl*cKO mice at 24 weeks old with the adjusted p-value <0.05. The x-axis indicates the number of genes in each category; **D)** Normalized relative expression of *Epas1* (*Hif1a)*, *Epor*, and *Slc2a1*(*Glut1)* from 10-week-old (circles) and 24-week-old (triangles) whole bone marrow of control or *Vhl*cKO mice.; **E)** *Epor* and **F)** *Slc2a1*(*Glut1)* normalized gene expression of 10-week-old (circles) and 24-week-old (triangles) sorted myeloid progenitors of control or *Vhl*cKO mice. * p < 0.05 ** p < 0.01, two-tailed Student’s t-test.

### Increased erythropoietin expression in the bone marrow of VhlcKO mice

To determine what may be driving the augmented RBC development in *Vhl*cKO mice, we measured EPO and TPO given that they act in synergy to expand the erythroblast populations^33^. EPO concentrations in the PB serum and BM fluid were comparable in both control and *Vhl*cKO mice at 3-to-6-weeks-old. However, EPO concentration was increased in the PB serum and BM fluid of 10-24-week-old *Vhl*cKO mice (**Figure 2D**). Interestingly, TPO concentration was increased only in the PB serum (**Figure 2D**).

To investigate if the elevated levels of EPO found in the *Vhl*cKO BM fluid were originating from BM cells, we measured *Epo* mRNA levels by real-time qPCR. Only one out of 4 control mice bone marrow had detectable, but very low, levels of *Epo (Ct value = 39.7)*, while the other 3 mice had no detectable levels after 40 cycles of qPCR. However, we detected lower delta cycle-threshold (C_t_) values of *Epo* mRNA within *Vhl*cKO BM cells, indicating a high expression of *Epo* mRNA (**Figure 2D**). We also measured *EpoR* mRNA expression from BM cells and found a decrease in *EpoR* relative expression in the *Vhl*cKO mice (**Figure 3D**).

### Elevated RBC parameters in the peripheral blood of VhlcKO mice

We expected the observed skewing towards erythropoiesis in the BM to be corroborated by hematological parameters in the peripheral blood. We found no significant changes in the WBC and differential counts at all timepoints, apart from the lymphocyte count being significantly decreased in the *Vhl*cKO at 6 weeks of age (**Supplemental Figure 4A-E**). However, as early at 6 weeks of age, we observed increased hemoglobin and hematocrit in *Vhl*cKO mice, and increased RBC counts starting at 10 weeks of age (**Figure 4A**), These changes were sustained to 24 weeks of age, and suggest *Vhl*cKO mice possess increased oxygen carrying capacity. Interestingly, by 10 weeks of age, *Vhl*cKO mice also exhibited increased mean corpuscular volume (MCV) and red blood cell distribution width (RDW) (**Supplemental Figure 4F, G**), corroborating the augmented erythroblast development observed in the BM. Additionally, at 24 weeks, *Vhl*cKO mice exhibited a significant decreased PLT count, but no significant changes in platelet volume or size (**Supplemental Figure 4H**). These findings demonstrate that *Vhl* deletion in *Dmp1*-expressing cells may not only be altering the state of erythropoiesis, but also the state of thrombopoiesis.

**Figure 4.**
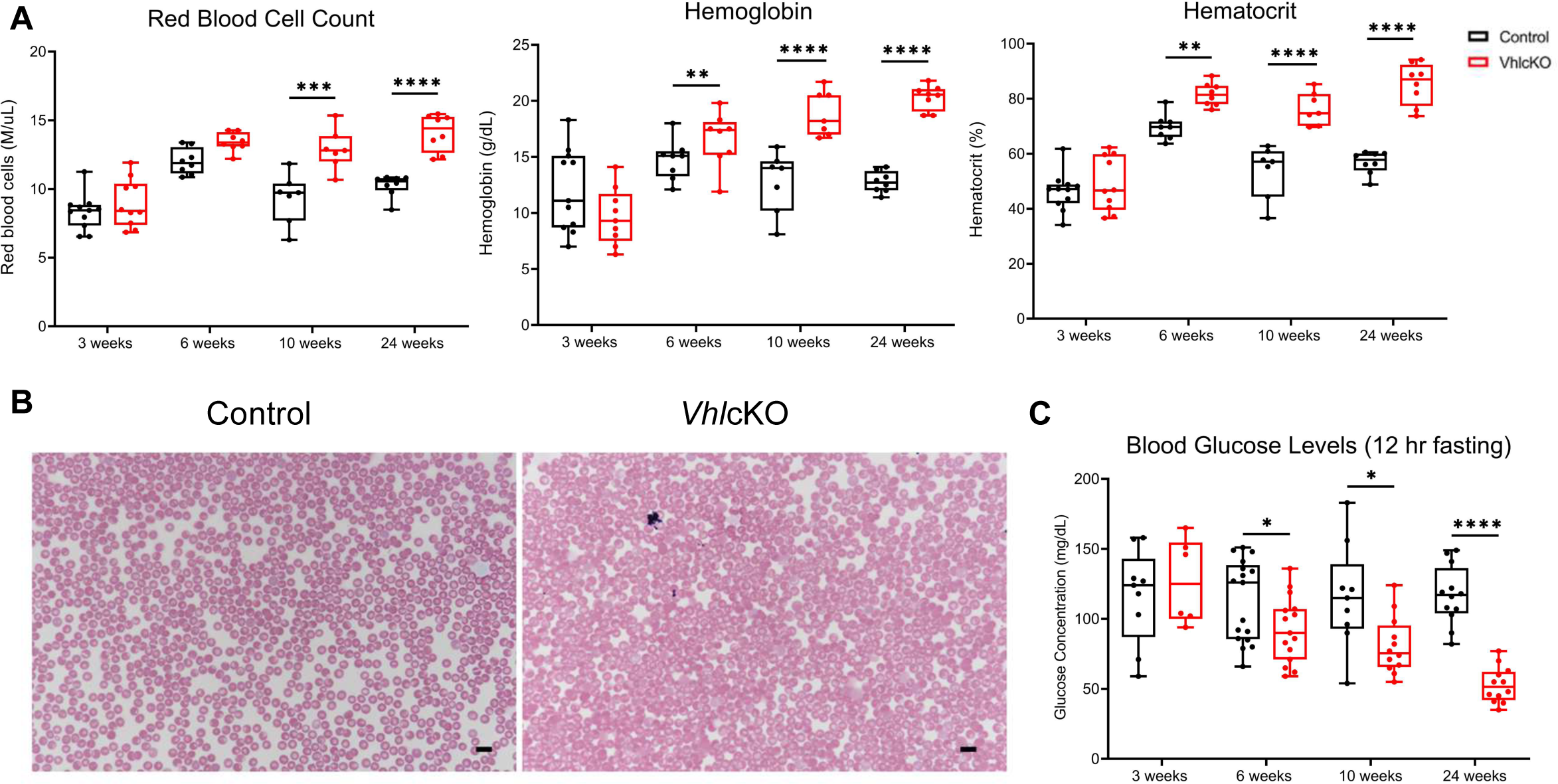
*Vhl*cKO mice have increased number of red blood cells with normal morphology but a low blood glucose level. **A)** Red blood cell count, hemoglobin, and hematocrit; **B)** Wright-Giemsa-stained peripheral blood smears of 10-week-old control and *Vhl*cKO mice, Scale bar: 10 µm; **C)** Blood glucose levels. Mice were fasted for 12 hours prior to glucose measurement. * p < 0.05 **** p < 0.0001, two-tailed Student’s t-test.

To examine the possibility of a hematological disorder, we performed PB smears as a follow-up to assess RBC morphology. In both control and *Vhl*cKO mice at 3, 6, and 10 weeks of age, the majority of RBC presented as spherical, anucleate biconcave discs with central pallor which is typical of healthy murine RBC^34^ (**Supplemental Figure 5A, Figure 4B**). We also observed marked polychromasia, which is a normal finding in mice; it is characterized by cytoplasmic basophilia resulting from immature RBC containing remnants of RNA^34^ (**Supplemental Figure 5A**). We attempted to perform PB smears on 24-week-old *Vhl*cKO mice, the blood was too viscous (likely because of high hematocrit (**Figure 4A**)), which prevented conclusive smearing to accurately assess RBC morphology.

### VhlcKO mice displayed a polycythemic phenotype

Polycythemia refers to increased RBC mass; this is reflected by an increase hemoglobin levels or hematocrit^35^. Our analysis of RBC parameters by CBC revealed that *Vhl*cKO mice have higher hematocrit and hemoglobin than control mice starting at 6 weeks of age (**Figure 4A**), implying a polycythemic phenotype is present. Our polycythemia diagnosis was further supported with our findings of no significant changes to WBC or PLT (**Supplemental Figure 4**). These findings suggest that only RBC overproduction is occurring, ruling out polycythemia vera, a subtype of polycythemia where overproduction of WBC and PLT also occurs. *Vhl*cKO mice also developed redness in their snouts and paws starting at 10 weeks of age (**Supplemental Figure 5B**), which may be a consequence of erythrocytosis. To further characterize what disorders may occur secondary to polycythemia, body weight and blood glucose levels were examined. Body weight was comparable between control and *Vhl*cKO mice (**Supplemental Figure 5C**), and starting at 6 weeks of age, *Vhl*cKO mice presented with hypoglycemia (**Figure 4C**). These findings suggest increased glucose consumption as a result of excess RBC in circulation^36^. Together, our data show that *Vhl* deletion in *Dmp1*-expressing cells results in polycythemia.

### Increased erythropoietic activity in spleens of VhlcKO mice

Previously, we found that *Vhl*cKO mice display splenomegaly (**Figure 5A**), characterized by increased spleen length, weight, and cellularity compared to controls^16^. While splenomegaly can be caused by increased splenic function, infiltration, or congestion^37^, we hypothesized that irregularities in the BM trigger the need for the spleen to increase its hematopoietic capacities. To test this hypothesis, we utilized CD71 and TER119 as cell surface markers to establish the state of erythropoiesis in the spleen (**Figure 5B**). Starting at 6 weeks of age, *Vhl*cKO spleens displayed increased frequency of BasoE and the presence of late-stage erythroblasts (**Figure 5C, 5D**). While cessation of splenic erythropoiesis typically occurs by 6 weeks of age in mice^38^, our findings suggest that changes in erythropoietic activity in the *Vhl*cKO spleen may be driven by irregularities in the BM^16^.

**Figure 5.**
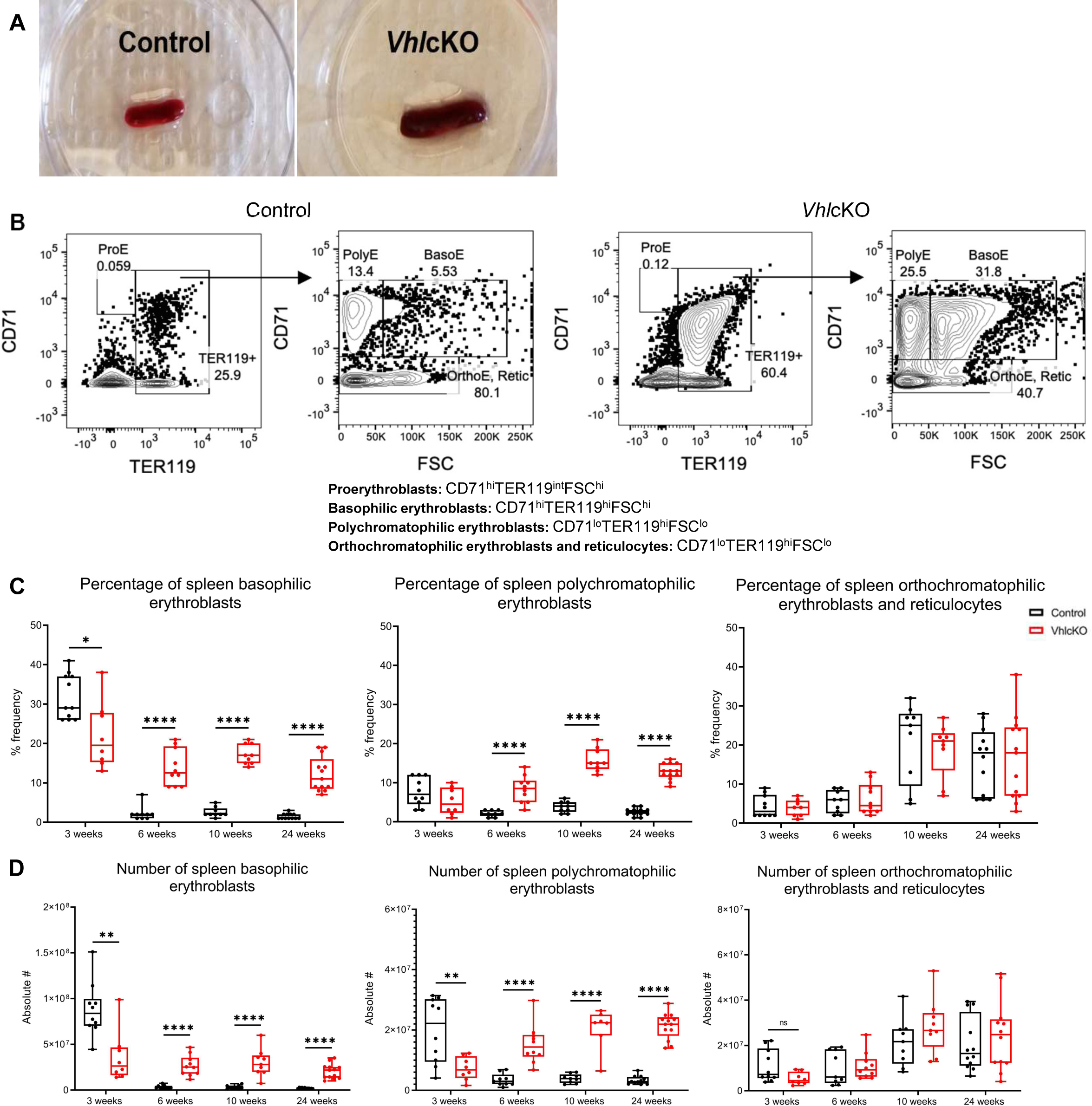
*Vhl*cKO mice have increased spleen erythropoietic activity. **A)** White light images of control spleen (left) and *Vhl*cKO spleen (right); **B)** Flow cytometry gating of erythroblast populations in the spleen of 24-week-old control and *Vhl*cKO mice. ProE, proerythroblasts; BasoE, basophilic erythroblasts; PolyE, polychromatophilic erythroblasts; OrthoE and Retic, orthochromatophilic erythroblasts and reticulocytes; **C)** Percentage and **D)** Number of erythroblast populations. * p < 0.05 ** p < 0.01 **** p < 0.0001, two-tailed Student’s t-test.

To determine if the observed splenomegaly was due to increased hematopoietic function and specific to the spleen, we performed hematoxylin and eosin staining of the spleens and livers of control and *Vhl*cKO mice. Starting at 6 weeks of age, we found significant red pulp hyperplasia and presence of megakaryocytes in the spleens of *Vhl*cKO mice, which became severe at 24 weeks of age (**Figure 6A** and **Supplemental Figure 6A).** While 3- and 6-week-old control and *Vhl*cKO livers were comparable, 10- and 24-week-old *Vhl*cKO livers exhibited hepatic hypertrophy and extramedullary hematopoiesis (**Supplemental Figure 6C**), as characterized by the many hepatocytes containing enlarged or multiple nuclei and small aggregates of erythroid cells with basophilic nuclei, respectively^39,40^. These findings show that *Vhl* deletion in *Dmp1*-expressing cells is inducing extramedullary hematopoiesis in both the spleen and liver to compensate for BM microenvironmental irregularities.

**Figure 6.**
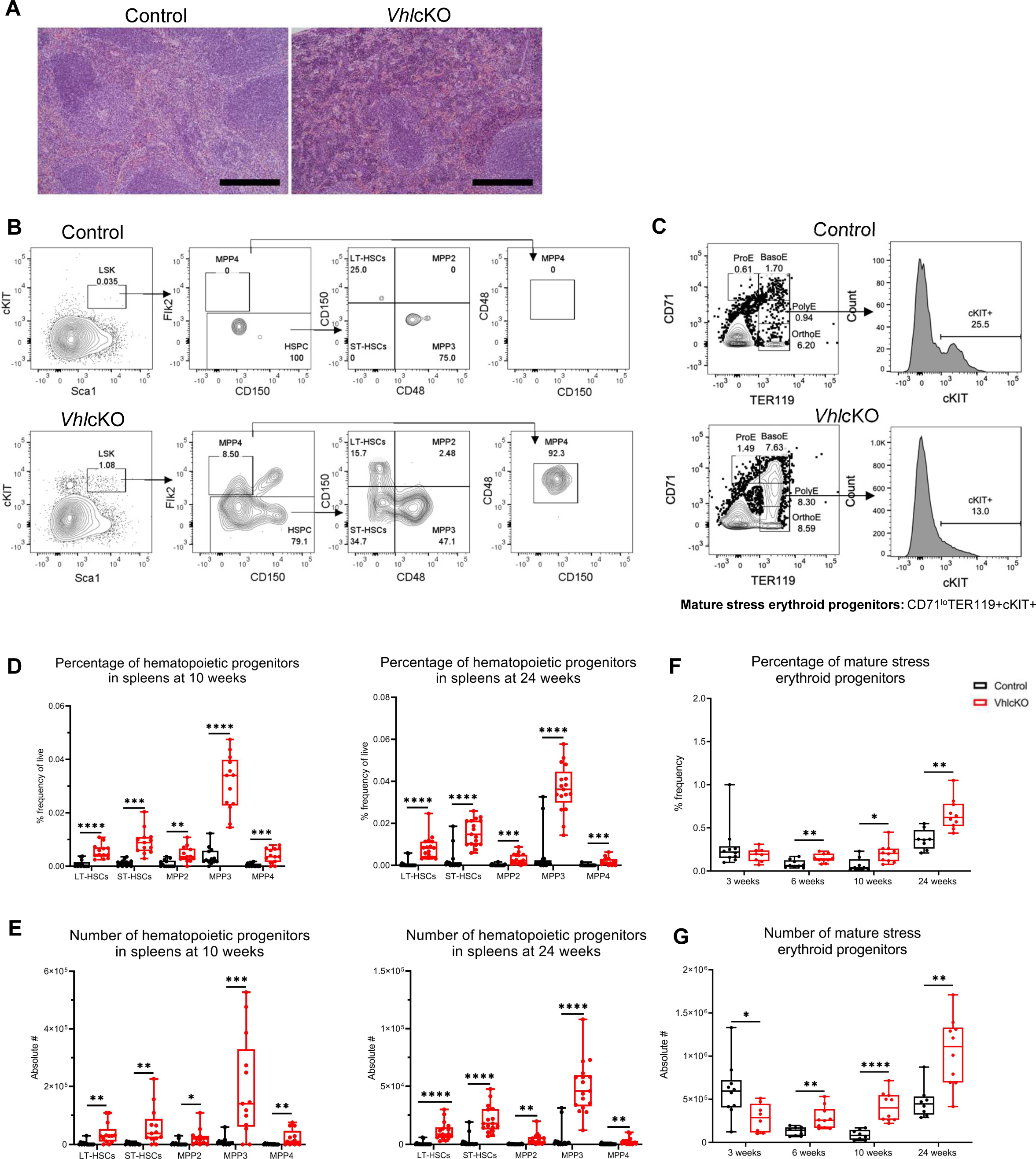
Evidence for extramedullary hematopoiesis and EPO-dependent stress erythropoiesis in the spleens of *Vhl*cKO mice. **A)** Photomicrographs of control spleen (left) and *Vhl*cKO spleen (right) at 24 weeks of age, Scale bar: 100 μm. Both groups display distinct regions of the red pulp (stained pink by eosin stain) and the white pulp (stained purple by hematoxylin); **(B)** Flow cytometry gating of hematopoietic progenitors in spleens control and *Vhl*cKO mice; **C)** Flow cytometry gating of the mature SEPs in the spleens of control and *Vhl*cKO mice; **D)** Percentage and **E)** Number of hematopoietic progenitors at 10 weeks (left panel) and 24 weeks (right panel); **F)** Percentage and **G)** Number of stress erythroid progenitors over time. * p < 0.05 ** p < 0.01 *** p < 0.001 **** p < 0.0001, two-tailed Student’s t-test.

### Evidence for EPO-dependent stress erythropoiesis in VhlcKO spleens

While it is known that developmental cues stimulate the spleen, conditions of stress (i.e. anemia) or underlying diseases (caused by conditions like nutrient deficiency) induce stress erythropoiesis as a mechanism to temporarily rescue steady state erythropoiesis^7^. This response is driven by SEPs, which are distinct from the progenitors involved in steady state erythropoiesis^41^. To determine if SEPs are present, we first enumerated hematopoietic progenitors in the spleen. We quantified long-term hematopoietic stem cells (LT-HSCs: LSK, CD150+ CD48-), short term hematopoietic stem cells (ST-HSCs: LSK, CD150-, CD48-), multipotent progenitors (MPP2: LSK, CD150+, CD48+; MPP3: LSK, CD150-, CD48+; and MPP4: LSK, CD150-, Flk2+, CD48+) (**Figure 6B)**. We found increased absolute number and frequency of all hematopoietic progenitors in the spleen of 10- and 24-week-old *Vhl*cKO mice (**Figure 6D-E**). Next, knowing that SEPs are derived from ST-HSCs that home to the spleen^7^, we enumerated mature SEPs (CD71^low^TER119^+^cKIT^+^) in the spleens of *Vhl*cKO mice using flow cytometry^7^ (**Figure 6C**). We found a lower absolute number of SEPs in 3-week-old *Vhl*cKO mice. However, starting at 6 weeks of age, we observed an increase in both the absolute number and frequency of SEPs in *Vhl*cKO mice (**Figure 6F-G**). Concurrently, we found that 10-to-24-week-old *Vhl*cKO mice exhibited increased EPO concentrations in their PB serum (**Figure 2A**), suggesting that EPO is driving the transition of SEPs to erythroblasts in the spleen^7,42^. These findings support the idea that EPO-dependent stress erythropoiesis is occurring because of BM irregularities in *Vhl*cKO mice.

## DISCUSSION

In the present study, we report that deletion of the *Vhl* gene in *Dmp1*-expressing cells results in hematopoietic cell-extrinsic changes in the BM microenvironment that alter the state of erythropoiesis as early as 6 weeks of age. We observed increased myeloid progenitor frequency, which suggests skewing towards myelopoiesis. We also observed upregulation of genes involved in response to glucose and actin filament arrangement in sorted MEPs, elevated EPO levels in both the PB serum and BM, and extramedullary hematopoiesis in the spleens and livers of *Vhl*cKO mice, all of which are consistent with augmented erythroblast development. To our knowledge, our report is the first to show the presence of mature SEPs *in vivo* due to *Vhl* deletion in bone, indicating that stress erythropoiesis may be occurring. In addition, our hematological analyses revealed a polycythemic phenotype in parallel to the changes observed in the bone. Altogether, these findings implicate HIF-driven alterations in skeletal homeostasis is driving changes in EPO levels and erythropoiesis, extending our insight to a functional relationship between these two tightly regulated processes.

Consistent with previous findings from our group and others^14–16^, our *Vhl*cKO mice displayed high bone mass, which can be attributed to the stabilization of HIF by conditionally deleting *Vhl* using the Cre/loxP system with *Dmp1* promoter driven Cre expression. In *VhlcKO* mice, *Vhl* deletion occurs in non-hematopoietic cells; therefore, any effects on hematopoietic cells observed must be cell extrinsic. Our RNA-seq data confirmed that *Vhl* is expressed in the MEPs of *Vhl*cKO mice, ruling out off-target expression of *Dmp1*-Cre in MEPs. In addition, *Vhl*cKO →wild-type whole BM chimeras displayed normal erythropoiesis (data not shown), further confirming that the hematopoietic progenitors developing in *Vhl*cKO mice are not skewed to differentiate into any particular lineage.

The molecular mechanisms and roles of VHL in facilitating cellular adaptation to hypoxic conditions via HIF-1 *α* and -2 *α* signaling are well-known. Functionally, HIF-1 *α* and - 2*α* heterodimerize with HIF-*β* in the nucleus of cells experiencing hypoxia to transcriptionally activate HIF target genes, including EPO to regulate erythropoiesis^43^. During development, the source of EPO gradually switches from the fetal liver to the kidney, which becomes the major site of EPO production during adulthood. To test for erroneous renal deletion of *Vhl,* we performed reverse transcription-PCR on 24-week-old *Vhl*cKO kidneys and found that *Vhl* expression was present, implying that kidney function is normal (**Supplemental Figure 7**). This suggests that in addition to local changes in *Vhl*cKO bones^12^, the increased EPO levels we observed in the BM fluid may also be driven by mechanisms involving the kidney^12,44,45^. While hypoxia and HIF signaling are the primary stimuli for renal EPO production, hepatocytes can also reactivate EPO expression in response to acute hypoxia^46^. Further experimentation is required to confirm the sources of EPO driving the changes in erythropoiesis in *Vhl*cKO mice.

With our finding of elevated EPO levels in both the PB serum and BM of our 10- and 24-week-old *Vhl*cKO mice, we expected to find local and systemic effects due to the pleiotropic nature of EPO. Flow cytometry analysis of BM erythroblast populations revealed increased number and frequency of OrthoE and Retic. Further supporting this data, our CBC analysis of RBC parameters revealed an increase in the number of RBC followed by increased levels of hematocrit and hemoglobin, suggesting that elevated EPO levels are inducing erythrocytosis in *Vhl*cKO mice. Moreover, *Vhl*cKO mice displayed splenomegaly with significant red pulp hyperplasia along with aggregates of erythroid cells within the liver, both of which are features of extramedullary hematopoiesis. These findings are consistent with previous reports using osteolineage-specific Cre drivers to conditionally delete *Vhl*^12,13^ and models studying polycythemia vera^47,48^. Unexpectedly, our CBC analysis also revealed increased MCV and RDW, indicating the RBC of *Vhl*cKO mice vary in size (anisocytosis)^34^. Although anisocytosis is a common finding in mice due to the degree of polychromasia present, it is also used as a differential diagnosis of anemia^49^. However, with erythrocytosis present, we suspect these data to correlate with the augmented erythroblast development in the BM. Typically, anisocytosis and erythrocytosis is associated with vascular abnormalities resulting in complications like thrombosis and hypertension, which inevitably can lead to premature lethality^50–52^. Intriguingly, our previous findings indicate that our *Vhl*cKO mice displayed significant vasodilated blood vessels in the BM^16^, suggesting that local and possibly systemic adaptive mechanisms are induced in these mice to compensate for the erythrocytosis. Such adaptive mechanisms may involve upregulated nitric oxide expression or reduced RBC lifespan to prevent highly viscous blood^48,53^. Further studies to evaluate these possibilities are necessary.

Given that alterations in erythropoiesis are present, we suspected that thrombopoiesis was also affected in *VhlcKO* mice. Our CBC analysis of platelet parameters revealed a decrease in the number of platelets in our 24-week-old *Vhl*cKO mice. Unexpectedly, we also observed elevated TPO levels in the peripheral blood serum starting at 3 weeks of age. Thrombopoietin is a glycoprotein that regulates megakaryopoiesis and thrombopoiesis; it is primarily produced by hepatocytes^54^. Hepatic mass and platelet count are directly linked, and clinical data shows that TPO mRNA expression is reduced in livers of patients with cirrhosis^55,56^. While decreased hepatic TPO production is a causative factor of thrombocytopenia in liver disease, we suspect our observations correlates with EMH or reflect dysregulation of platelet sequestration^57^. In this light, it seems plausible that increased platelet sequestration may be occurring in the spleen and contributing to the splenomegaly in our mice^58^.

We hypothesize that splenomegaly in our *Vhl*cKO mice occurs because of increased hematopoietic capacity to compensate for irregularities present in the BM microenvironment. Extensive analysis of anemic stress in mice has been characterized by increased splenic erythropoiesis to compensate for low BM erythroid output. However, we observed alterations in myelopoiesis, characterized by increased percentage of CMPs and MEPs present. These upstream changes indicate that HIF-driven changes in skeletal homeostasis may be contributing to the elevated EPO levels we observe, and thereby inducing reactivation of splenic erythropoietic properties in a novel manner. Our flow cytometry analysis of spleen erythroblast populations revealed the presence of late-stage erythroblasts. In support of this data, histological analysis of 10- and 24-week-old *Vhl*cKO spleens displayed extramedullary hematopoiesis that is primarily myeloid driven as seen with the presence of megakaryocytes and red pulp hyperplasia. This may indicate sequestration, accumulation, and proliferation of circulating myeloid progenitors in splenic cords is occurring. Following this thought, we presume this may also be due to the homing of ST-HSCs to the spleen, where the stress erythroid fate promotes the extramedullary nature of stress erythropoiesis^59^.

Unlike steady state erythropoiesis, which produces RBC at a constant rate, stress erythropoiesis generates a bolus of new RBC derived from SEPs. Early *in vivo* work indicated that these splenic progenitors were distinct from steady state erythroid progenitors^10^. Our data now provide *in vivo* evidence that terminal differentiation of SEPs may contribute to the red pulp hyperplasia and increased number of circulating RBC we observe in *Vhl*cKO mice. Alternatively, our data could reflect increased RBC turnover or sequestration due to erythrocytosis. We next considered the possibility that the elevated EPO levels in our mice are mimicking EPO stress. In contrary to studies analyzing recovery from anemia via EPO treatment, researchers also use EPO treatment to induce stress erythropoiesis^60^. Inducing stress erythropoiesis in this way causes the differentiation of erythroblasts and skews hematopoiesis to favor erythropoiesis, which might suggest that stress erythropoiesis could be considered EPO-dependent. While elevated EPO levels due to renal HIF stabilization is likely not occurring, further studies to evaluate oxygen and iron levels is necessary.

While initial studies have shown that activation of HIF signaling in osteolineage cells is anabolic to bone, early and current work support the idea that osteogenic cells are the main consumers of glucose in the skeleton^13,61,62^. In agreement with this, we observed a marked hypoglycemic phenotype in our *Vhl*cKO mice starting at 6 weeks of age. It is plausible that the increased number of RBC and their consequent uptake of glucose may be improving glucose clearance^63^. With this hypoglycemic phenotype occurring independent to changes in body weight, sex-dependent and hormonal changes need to be further explored^64^. Although we have not directly measured glucose tolerance, we hypothesize body fat and insulin sensitivity would be the main contributing factors to changes in glycemic control, as other studies have found^13,65^.

Our findings may have important implications for improving bone anabolic therapies for osteoporosis and potentially can serve as a therapeutic for individuals with anemia, particularly those with frank diabetes mellitus (DM) suffering from anemia of renal disease. While the etiology and clinical manifestations of osteoporosis, anemia, and DM are driven by varying but overlapping risk factors and lifestyle choices, often, these disorders occur as comorbidities^66–68^. Osteoporosis is diagnosed based on low bone mineral density (BMD) in addition to a low body mass index and increased risk of fracture^68^. Low BMD is often dismissed until complications (i.e., fracture) occur, leading to a marked decrease in quality of life with increased morbidity, mortality, and disability. As with osteoporosis, anemia may go underdiagnosed or unrecognized until clinical symptoms are present. For these reasons, efforts are focused on developing predictive factors for fracture risk and targeted treatments. In recent years, recombinant human erythropoietin (rHuEPO), a erythropoietin-stimulating agent (ESA) has become the standard therapy for treatment of anemia and has been shown to be a suitable alternative for glycemic control in patients with DM^69^. However, this kind of ESA therapy is expensive and requires a doctor’s visit for administration. Fortunately, PHD inhibitors and HIF modulators have emerged as an appealing therapeutic strategy to activate EPO under normoxia^70,71^. It is thought that these pharmacological therapies will not only revolutionize the treatment of anemia, but also the treatment of osteoporosis given its shared mechanism of action. Yet, it remains a challenge to target the HIF signaling pathway without dysregulating the skeletal and hematopoietic system^16,72,73^ and causing off target metabolic, hormonal, and renal effects^74,75^. Therefore, in the future, the benefits of tissue-specific inhibition of HIF signaling may help advance the paradigm of managing an isolated diagnosis or comorbidities of osteoporosis, diabetes, and anemia.

The findings in this study expands our understanding of the role of *Vhl* and altered skeletal homeostasis has on the state of erythropoiesis. Our data suggest that deleting *Vhl* in *Dmp1*-expressing MSCs, osteoblasts, and osteocytes induces HIF-driven alterations that not only causes BM microenvironmental irregularities, but also directly regulates the state of erythropoiesis through an increase in EPO and expansion of the erythroid niche to the spleen and liver (**Figure 7**). We suspect these changes in erythropoiesis are mediated mainly through systemic EPO secretion with local EPO secretion also contributing. Future studies investigating both the hepatic and splenic stress erythroid niche could provide insight as to how EPO and its co-regulators (i.e., FGF23) regulate the state of erythropoiesis to compensate for BM irregularities^76^. Nevertheless, these findings emphasize that when designing treatments for osteoporosis and anemia to select the most appropriate point of therapeutic intervention to modulate HIF activity among the proper dosage and route of administration^71^. Considering these factors may help to avoid unwanted side effects caused by activating undesired HIF target genes.

**Figure 7.**
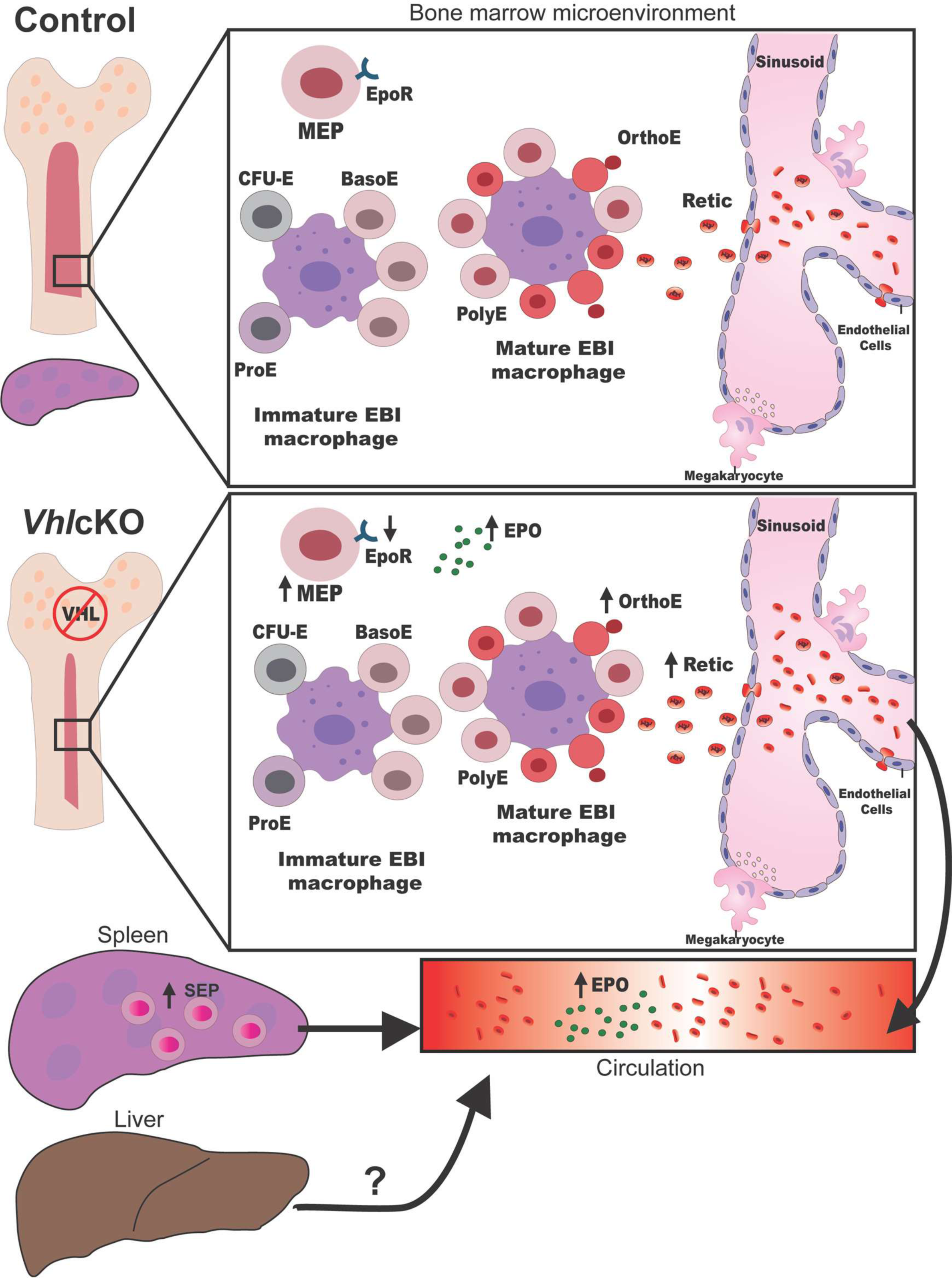
*Vhl* deletion in Dmp1-expressing cells contributes to alterations in the state of erythropoiesis. Schematic model describing the effects of BM microenvironmental changes on erythropoiesis extending from the bone marrow to periphery and spleen.

## ACKNOWLEDGMENTS

We thank the staff of the Department of Animal Research Services (DARS) at UC Merced for excellent animal care. We acknowledge Dr. David Gravano in the Stem Cell Instrumentation Foundry for assistance in generating flow cytometry data. Research was supported in part by the DoD Research and Education Program for HBCU/MSI Instrumentation Grant (W911NF1910529), the Genomics Research and Technology Hub Shared Resource of the Cancer Center Support Grant (P30CA-062203), the Single Cell Analysis Core shared resource of Complexity, Cooperation and Community in Cancer (U54CA217378), the Genomics-Bioinformatics Core of the Skin Biology Resource Based Center @ UCI (P30AR075047) at the University of California, Irvine and NIH shared instrumentation grants 1S10RR025496-01, 1S10OD010794-01, and 1S10OD021718-01. We also thank Dr. Katrina Hoyer, Dr. Anh Diep, and Christi Turner for access to and assistance with the Hemavet complete blood cell counting equipment. Part of the work was performed under the auspices of the US Department of Energy by Lawrence Livermore National Laboratory under contract no DE-AC52-07NA27344.

## AUTHOR CONTRIBUTIONS

JME, BC, CD and JOM designed the study, collected data, performed data analysis, and interpreted the results. HT, CP, and HP assisted in data acquisition. AS and BC analyzed the bulk RNA-seq data, SLT performed the histological analysis. JME, BC and JOM wrote the manuscript.

