## Supplemental Materials for "Vhl deletion in *Dmp1*-expressing cells alters MEP metabolism and promotes stress erythropoiesis"

**Supplemental Figure 1. Histology of sternum with bone marrow as a function of age.** Photomicrographs of control and *Vhl*cKO sternum at 3, 6, and 10 weeks of age, where bone is light pink colored and bone marrow is dark purple colored; scale bar: 100 μm.

**Supplemental Figure 2. *Vhl*cKO bone marrow myeloid lineage cells.** Percentage and number of **A)** Neutrophils**, B)** Dendritic cells and **C)** Monocytes from bone marrow of 3, 6, and 10-week-old control and *Vhl*cKO mice. * $p < 0.05$ ** $p < 0.01$ *** $p < 0.001$, two-tailed Student's t-test.

**Supplemental Figure 3. Flow cytometry gating strategy for myeloid progenitor sorting.** Example of myeloid progenitor gating strategy for sorting GMP, MEP and CMP from control (top) and *Vhl*cKO (bottom) pooled mice for qPCR and bulk RNA-Seq analysis. Plots from pre-sorted and post-sorted samples are shown.

**Supplemental Figure 4. *Vhl*cKO mice display changes in red blood cell volume and decreased platelets, but normal white blood cell differential. A)** White blood cell count; **B)** Neutrophil and lymphocyte count; **C)** Monocyte, eosinophil, and basophil count; **D)** Percentage of neutrophils, lymphocytes, and monocytes; **E)** Percentage of eosinophils and basophils; **F)** Mean corpuscular volume and mean corpuscular hemoglobin; **G)** Mean corpuscular hemoglobin concentration and red blood cell distribution width; **H)** Platelet count and mean platelet volume. * $p < 0.05$ ** $p < 0.01$ *** $p < 0.001$, two-tailed Student's t-test.

**Supplemental Figure 5. *Vhl*cKO mice have normal red blood cell morphology and percentage of polychromasia but present a polycythemic phenotype. A**) Wright-Giemsa-stained peripheral blood smears of control and *Vhl*cKO mice at 3 and 6 weeks of age, Scale bar: 10 μm; **B)** Starting at 10 weeks of age, the snout and paws of *Vhl*cKO mice appear redder in color than control mice. Ten-week-old mice are pictured. **C)** Body weight; * $p < 0.05$ **** $p < 0.0001$, two-tailed Student's t-test.

**Supplemental Figure 6. Evidence for extramedullary hematopoiesis in *Vhl*cKO mice. A)** Photomicrographs of control and *Vhl*cKO spleen at 3, 6, and 10 weeks of age, Scale bar: 100μm. Both groups display distinct regions of the red pulp (stained pink by eosin stain) and the white pulp (stained purple by hematoxylin). **B)** Percentage and number of spleen myeloid progenitors from 10- and 24-week-old control and *Vhl*cKO mice; **C)** Photomicrographs of control and *Vhl*cKO livers at 3, 6, 10, and 24 weeks of age; Scale bar: 100μm. * $p < 0.05$ *** $p < 0.001$ **** $p < 0.0001$, two-tailed Student's t-test.

**Supplemental Figure 7. Validation of *Vhl* expression in *Vhl*cKO kidneys.** Reverse transcription-PCR validation of *Vhl* and *Gapdh* expression in whole kidney cells of 24-week-old control (lanes 1-3) and *Vhl*cKO (lanes 4-6) mice. Abbreviations: Kid, kidney; -RT, no reverse transcriptase.

**Supplemental Table 1:** List of the fluorochrome-labeled monoclonal antibodies used for flow cytometry by experimental cocktail

**Supplemental Table 2:** List of differentially expressed genes (DEG) from sorted MEPs of VhlcKO and control mice

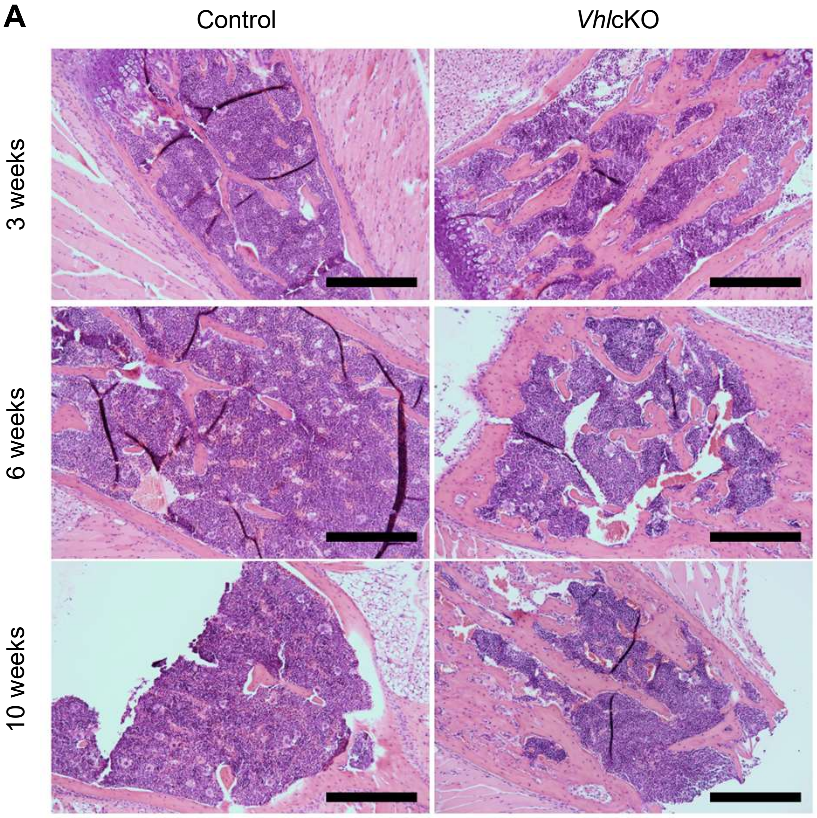


Supp Figure 1

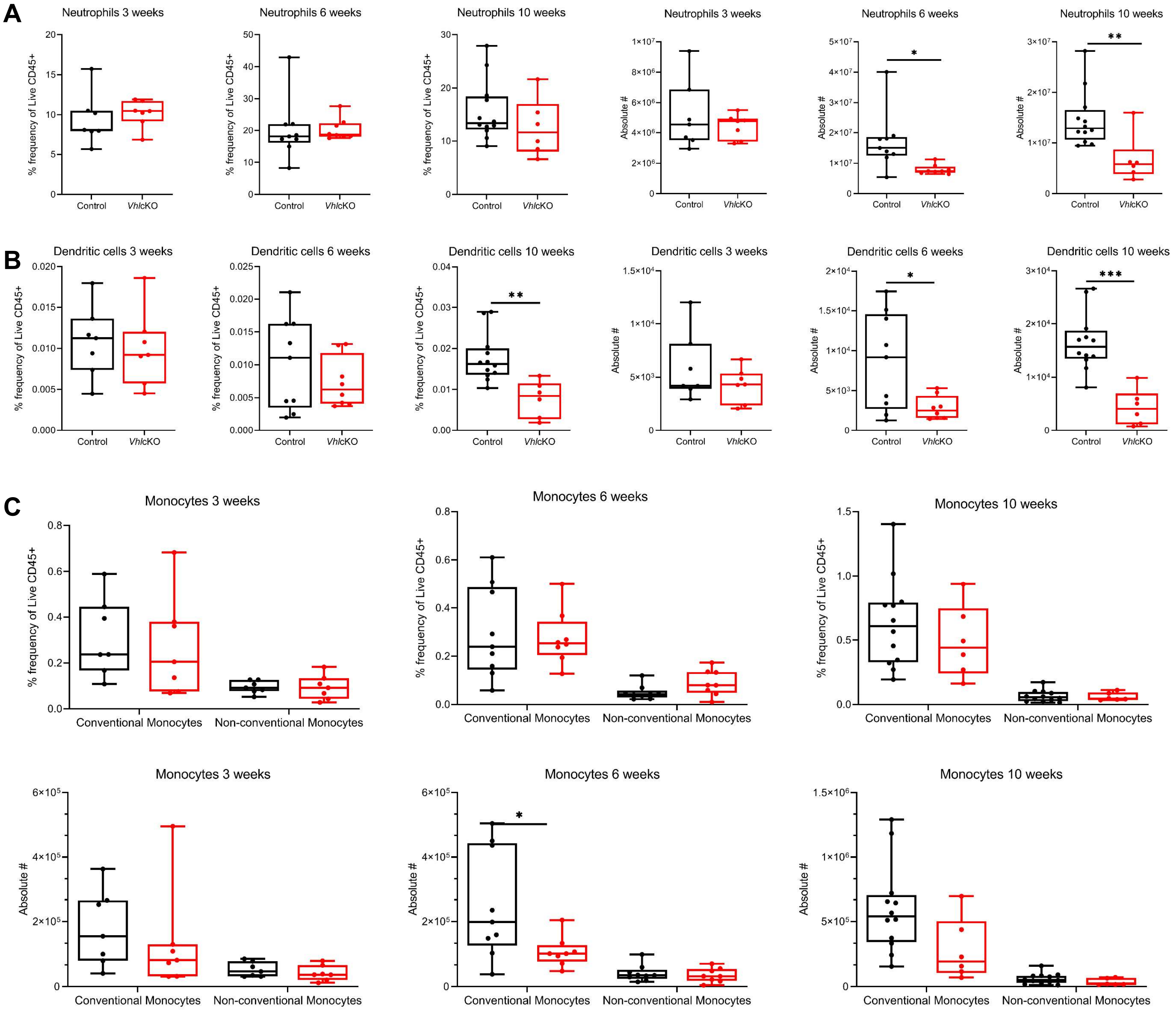


Supp Figure 2

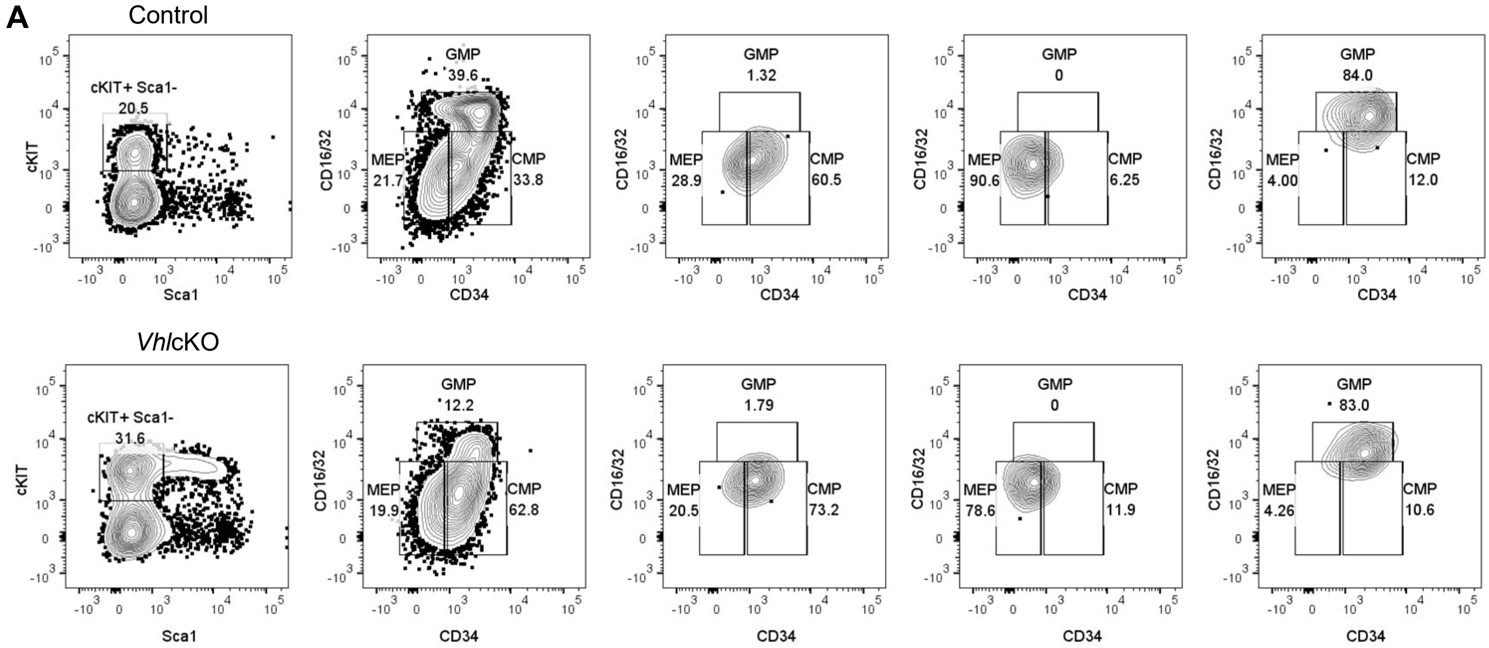


Supp Figure 3

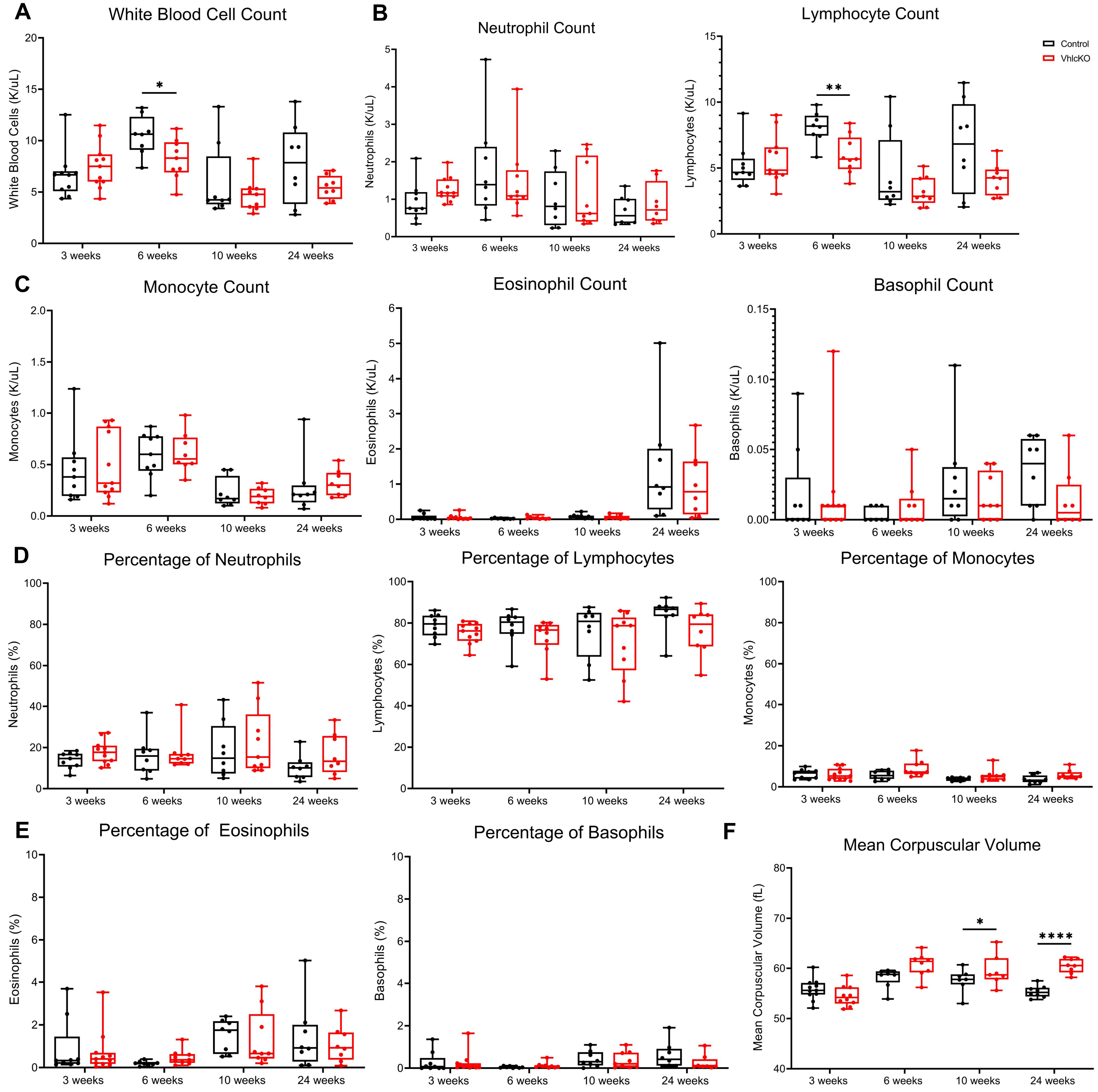


Supp Figure 4

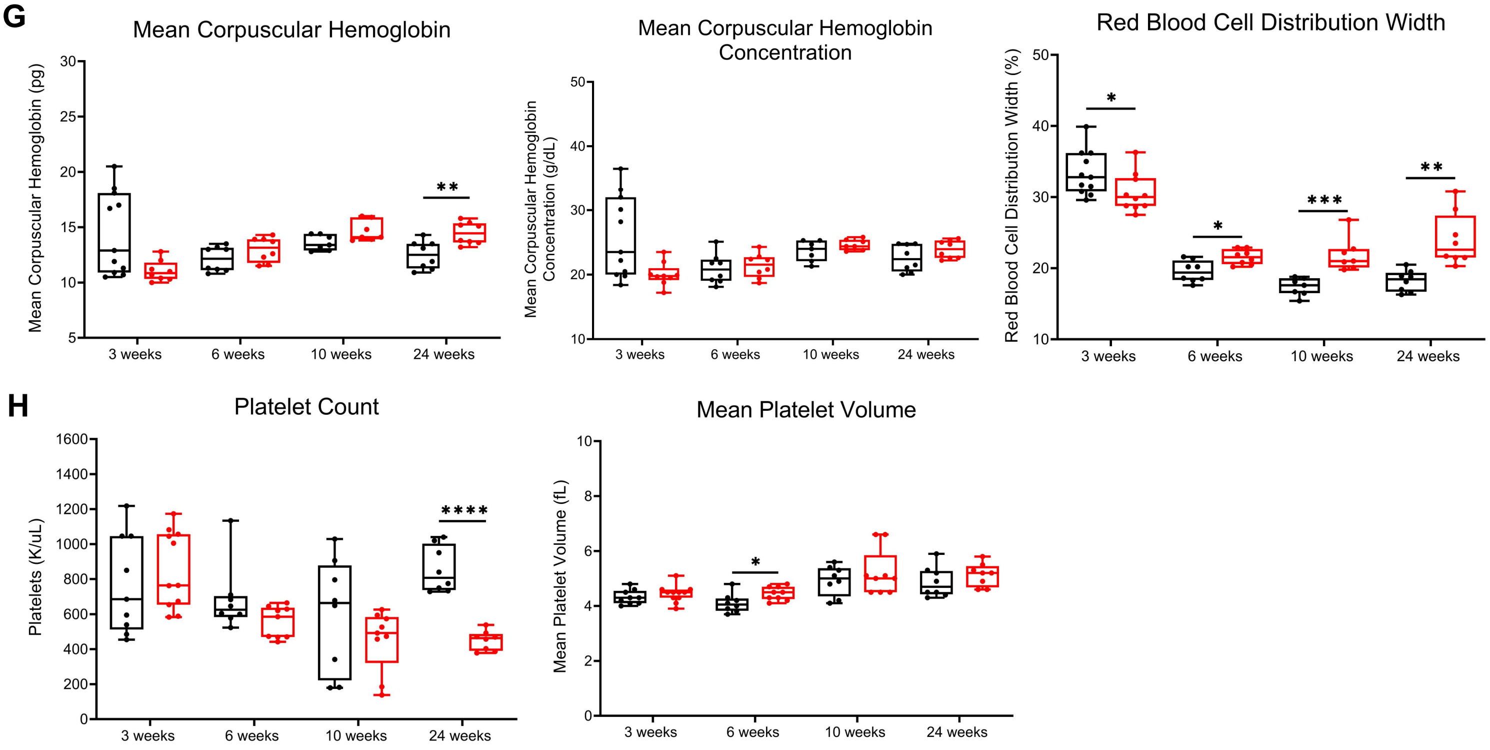


Supp Figure 4 continued

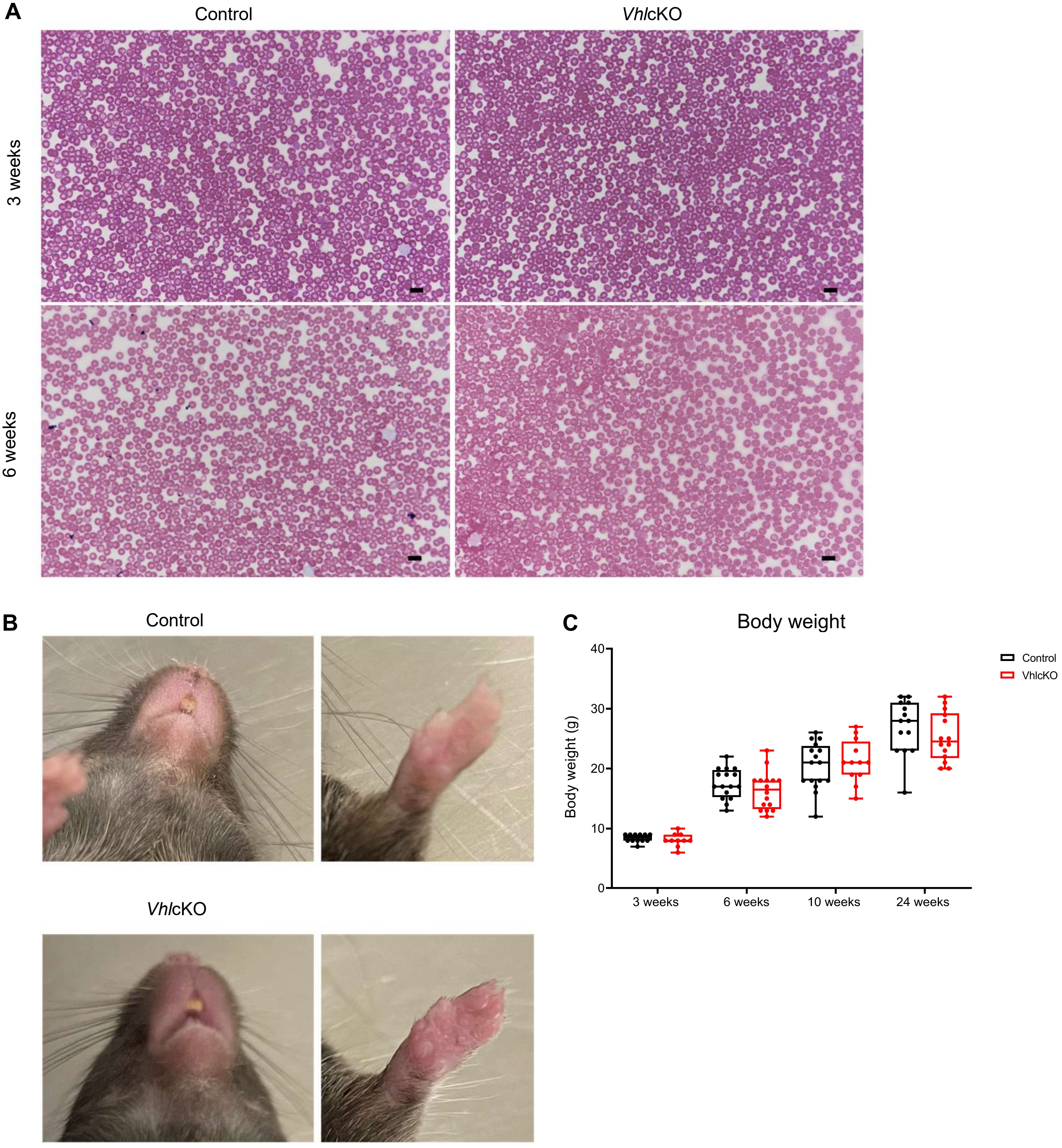


Supp Figure 5

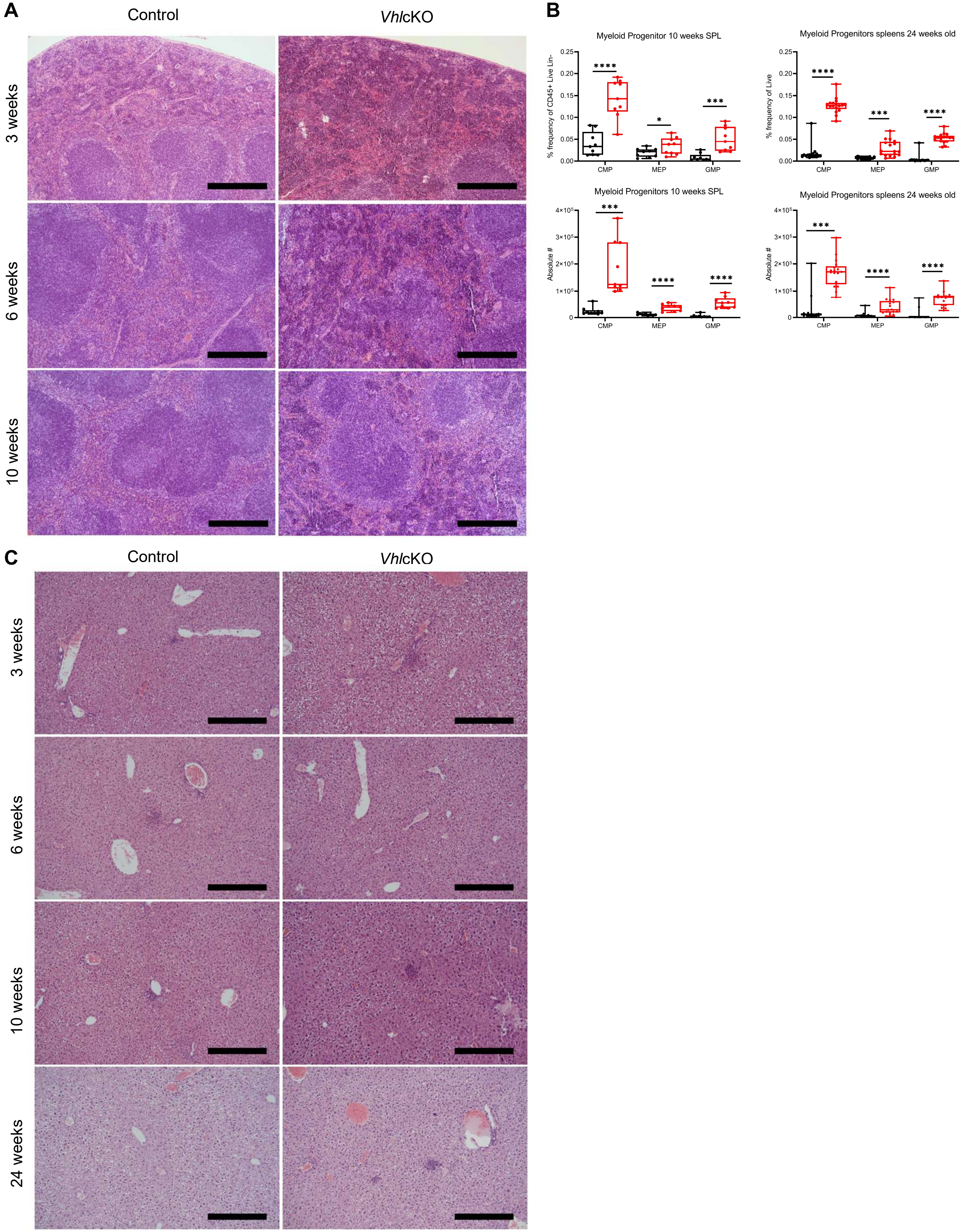


Supp Figure 6

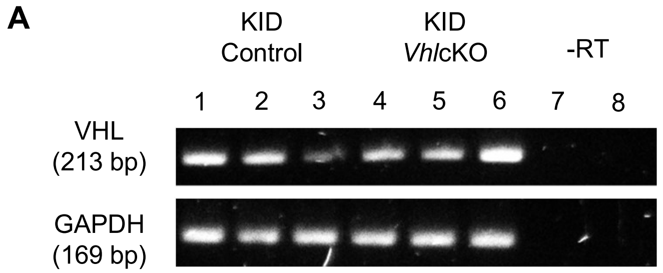


Supp Figure 7

**Supplemental Table 1.** List of the fluorochrome-labeled monoclonal antibodies used for flow cytometry by experimental cocktail

| Cocktail | Antigen | Clone | Fluorochrome | Source | Target population markers |
|---|---|---|---|---|---|
| Myeloid Lineage | CD11b | M1/70 | FITC | Biolegend | neutrophils: CD45$^{+}$Ly6G$^{+}$CD11b$^{+}$ |
| | CD11c | N418 | PE | Biolegend | |
| | CD19 | 6D5 | BV421 | Biolegend | |
| | CD43 | S7 | BV605 | BD Biosciences | classical monocytes: CD45$^{+}$Ly6G$^{-}$CD11b$^{+}$F4/80$^{-}$ CD115$^{+}$Ly6C$^{hi}$ |
| | CD45 | 30-F11 | BV711 | Biolegend | |
| | Ly6G | 1A8 | APC | Biolegend | |
| | F4/80 | BM8 | APC-eFluor 780 | Invitrogen | non-classical monocytes: CD45$^{+}$Ly6G$^{-}$ CD11b$^{+}$F4/80$^{-}$CD115$^{+}$Ly6C$^{-}$CD43$^{+}$ |
| | Ly6C | HK1.4 | PerCP Cy5.5 | Biolegend | |
| | CD115 | AFS98 | PE-Cy7 | Invitrogen | |
| | MHC-II | M5/114.15.2 | biotin | Biolegend | dendritic cells: CD45$^{+}$Ly6G$^{-}$CD11b$^{+}$F4/80$^{-}$ CD115$^{+}$Ly6C$^{-}$CD43$^{+}$MHC-II$^{+}$CD11c$^{+}$ |
| | Streptavidin | - | BUV395 | BD Biosciences | |
| | DAPI (4′,6-diamidino-2-phenylindole) | | Viability Dye | Sigma-Aldrich | |
| Myeloid Progenitor | CD3 | 145-2C11 | biotin | Biolegend | |
| | CD4 | Gk1.5 | biotin | Biolegend | |
| | CD8 | 53.6.7 | biotin | Biolegend | common myeloid progenitor: CD45$^{+}$Lin$^{-}$ cKIT$^{+}$Sca1$^{-}$CD16/32$^{+}$CD34$^{+}$ |
| | CD5 | 52-7.3 | biotin | Biolegend | |
| | CD19 | 6D5 | biotin | Biolegend | |
| | Nk1.1 | PK136 | biotin | Biolegend | |
| | TER119 | TER119 | biotin | Biolegend | |
| | Gr1 | RB6-8C5 | biotin | Biolegend | |
| | F4/80 | BM8 | biotin | Biolegend | granulocyte/monocyte progenitors: CD45$^{+}$Lin$^{-}$ cKIT$^{+}$Sca1$^{-}$CD16/32$^{-}$CD34$^{+}$ |
| | CD34 | RAM34 | FITC | Invitrogen | |
| | CD45 | 30-F11 | BV421 | Biolegend | |
| | Sca1 (Ly-6A/E) | D7 | BV510 | Biolegend | |
| | CD16/32 | 93 | APC | Biolegend | |
| | CD117 (cKit) | 2B8 | APC-Cy7 | Biolegend | megakaryocyte/erythroid progenitors: Lin$^{-}$ CD45$^{+}$cKIT$^{+}$Sca1$^{-}$CD16/32$^{-}$CD34$^{-}$ |
| | Streptavidin | - | PE-Cy5 | Life Technologies | |
| | Propidium iodide (PI) | | Viability Dye | Sigma-Aldrich | |
| Red Blood Cell Maturation | CD71 | RI7217 | PE | Biolegend | proerythroblasts: CD71$^{hi}$TER119$^{int}$FSC$^{hi}$ |
| | TER119 | TER119 | APC | Biolegend | basophilic erythroblasts: CD71$^{hi}$TER119$^{hi}$FSC$^{hi}$ |
| Stress Erythroid Progenitor | CD71 | RI7217 | PE | Biolegend | polychromatophilic erythroblasts: CD71$^{lo}$TER119$^{hi}$FSC$^{lo}$ |
| | TER119 | TER119 | APC | Biolegend | orthochromatophilic erythroblasts and reticulocytes: CD71$^{lo}$TER119$^{hi}$FSC$^{lo}$ |
| | CD117 (cKit) | 2B8 | APC-Cy7 | Biolegend | |

**Supplemental Table 2.** List of differentially expressed genes (DEG) from sorted MEPs of *VhlcKO* and control mice

| Gene | logFC_KOvsWT | P.Value | adj.P.Val |
|---|---|---|---|
| Lgals4 | 3.674673036 | 1.07E-05 | 0.045742556 |
| Podxl | 3.318478294 | 2.66E-05 | 0.045742556 |
| Clca1 | 5.158214665 | 3.12E-05 | 0.045742556 |
| Pigr | 5.246187763 | 3.51E-05 | 0.045742556 |
| Lypd8 | 5.501162003 | 3.88E-05 | 0.045742556 |
| Nr4a1 | 2.570935471 | 4.22E-05 | 0.045742556 |
| Krt8 | 4.967620955 | 4.47E-05 | 0.045742556 |
| Cyp2r1 | 3.98280926 | 7.06E-05 | 0.045742556 |
| Lgals3 | 3.541184788 | 8.07E-05 | 0.045742556 |
| Atp8b1 | 2.872147298 | 8.56E-05 | 0.045742556 |
| Dmbt1 | 4.900652648 | 9.70E-05 | 0.045742556 |
| Cobl | 6.393989041 | 0.00010645 | 0.045742556 |
| Satb2 | 2.787662435 | 0.00011178 | 0.045742556 |
| 2610528A11Rik | 6.909129751 | 0.00011467 | 0.045742556 |
| Klf4 | 2.028655708 | 0.00011475 | 0.045742556 |
| Mgat3 | 6.758190579 | 0.00012154 | 0.045742556 |
| Hmgcs2 | 4.055936503 | 0.00012555 | 0.045742556 |
| Tead1 | 6.538002803 | 0.00013284 | 0.045742556 |
| Aoc1 | 6.743280912 | 0.00013723 | 0.045742556 |
| Hnf4a | 3.634025833 | 0.00014067 | 0.045742556 |
| App | 2.1226055 | 0.00014302 | 0.045742556 |
| Mep1a | 6.713731871 | 0.00015348 | 0.045742556 |
| Spink4 | 3.648083366 | 0.00015955 | 0.045742556 |
| Tmprss2 | 5.483664641 | 0.0001644 | 0.045742556 |
| Ceacam2 | 3.323574501 | 0.00016726 | 0.045742556 |
| Clmn | 5.761073109 | 0.00017278 | 0.045742556 |
| Fcgbp | 5.517965839 | 0.00017611 | 0.045742556 |
| Abcc3 | 5.193607972 | 0.00017921 | 0.045742556 |
| Sema4g | 3.439889994 | 0.00020056 | 0.048998459 |
| Wwc1 | 6.529551962 | 0.00021215 | 0.048998459 |
| Sdcbp2 | 5.997955107 | 0.00021246 | 0.048998459 |
| Mgat4c | 5.740263634 | 0.0002133 | 0.048998459 |
| Krt19 | 5.622404321 | 0.00023598 | 0.051363713 |
| Phgr1 | 7.219735568 | 0.00023701 | 0.051363713 |
| Robo3 | 2.415260361 | 0.00024694 | 0.051781677 |
| Duox2 | 5.898628244 | 0.00025071 | 0.051781677 |
| Myh14 | 4.652098463 | 0.00025693 | 0.051781677 |
| Fermt1 | 5.32861827 | 0.00025697 | 0.051781677 |
| Myo1a | 4.252809221 | 0.00027317 | 0.053179672 |
| Fat1 | 4.961551356 | 0.00029118 | 0.054158455 |

| | | | |
|---|---|---|---|
| B4galnt2 | 2.410718809 | 0.00030008 | 0.054158455 |
| Ifit1bl1 | 5.774146851 | 0.00031004 | 0.054158455 |
| Oit1 | 5.439041616 | 0.00031028 | 0.054158455 |
| Shroom3 | 4.261886945 | 0.00031563 | 0.054158455 |
| Tnfrsf11a | 6.009833477 | 0.00031592 | 0.054158455 |
| Capn13 | 5.594288117 | 0.00032565 | 0.054338128 |
| Nov | 6.101246437 | 0.00034152 | 0.054578055 |
| Chp2 | 6.028864768 | 0.00034462 | 0.054578055 |
| Ceacam20 | 6.187271178 | 0.00035593 | 0.054578055 |
| Cdhr2 | 4.689059867 | 0.00035627 | 0.054578055 |
| Ptprf | 3.420266495 | 0.00036005 | 0.054578055 |
| Prkaa2 | 5.104158192 | 0.00036471 | 0.054578055 |
| Apol10a | 5.016316256 | 0.0003663 | 0.054578055 |
| Epcam | 5.292321359 | 0.00037024 | 0.054578055 |
| Il13ra1 | 5.980251213 | 0.00037063 | 0.054578055 |
| Guca2a | 4.537439872 | 0.00038509 | 0.055940012 |
| Exph5 | 6.266147703 | 0.0003985 | 0.055940012 |
| Sik1 | 1.291288162 | 0.00040031 | 0.055940012 |
| Klk1 | 5.828117334 | 0.00040319 | 0.055940012 |
| Reg3b | 5.468283636 | 0.00040423 | 0.055940012 |
| Abcb1a | 4.535630435 | 0.00041207 | 0.056346137 |
| Mtus1 | 4.473239056 | 0.0004245 | 0.056654814 |
| Nfib | 6.078723016 | 0.00042913 | 0.056654814 |
| Vil1 | 3.608215548 | 0.00044857 | 0.058388308 |
| S100a6 | 4.666394758 | 0.0004568 | 0.058388308 |
| Pck1 | 6.677329737 | 0.00045803 | 0.058388308 |
| Gpa33 | 5.263392525 | 0.00046387 | 0.058388308 |
| Myo7b | 5.248633987 | 0.00047188 | 0.058388308 |
| Tmprss4 | 4.879850808 | 0.00047731 | 0.058388308 |
| Emcn | 5.73350997 | 0.00047784 | 0.058388308 |
| Hsd3b3 | 6.109215692 | 0.00048377 | 0.058490547 |
| Akr1c14 | 6.482081013 | 0.00049971 | 0.058605381 |
| Cdh1 | 4.496868119 | 0.00050145 | 0.058605381 |
| Epb41l3 | 6.087712676 | 0.00050399 | 0.058605381 |
| Sectm1b | 5.230473437 | 0.00050513 | 0.058605381 |
| Slc4a4 | 5.362782753 | 0.00051855 | 0.059371202 |
| Irf2bpl | 1.774492058 | 0.00052207 | 0.059371202 |
| Krt18 | 2.441123249 | 0.00052775 | 0.059428936 |
| Emp1 | 4.143392011 | 0.00054191 | 0.059654409 |
| Jup | 2.731529209 | 0.00054989 | 0.059654409 |
| Reg4 | 5.663386849 | 0.00055336 | 0.059654409 |
| Esrp1 | 4.971668974 | 0.0005566 | 0.059654409 |
| Tff3 | 6.261086122 | 0.00056012 | 0.059654409 |

| | | | |
|---|---|---|---|
| Hist1h4m | -1.092263178 | 0.00057977 | 0.059661934 |
| Tspan1 | 5.063973745 | 0.00058012 | 0.059661934 |
| Aldh1b1 | 3.073641184 | 0.00058398 | 0.059661934 |
| Lima1 | 4.633792164 | 0.00061403 | 0.059661934 |
| Tfcp2l1 | 4.180166398 | 0.0006145 | 0.059661934 |
| Myo5b | 3.333390993 | 0.00061632 | 0.059661934 |
| Pard3 | 4.78775124 | 0.00061667 | 0.059661934 |
| Atp1b1 | 4.109514687 | 0.00062133 | 0.059661934 |
| Elf3 | 4.364598868 | 0.00062922 | 0.059661934 |
| Areg | 5.308494846 | 0.00063435 | 0.059661934 |
| Erbb2 | 3.896106631 | 0.00064023 | 0.059661934 |
| Rasef | 6.249046037 | 0.0006495 | 0.059661934 |
| Myo5c | 4.465126215 | 0.00064984 | 0.059661934 |
| Klf5 | 2.243825833 | 0.00065098 | 0.059661934 |
| Boc | 5.944643866 | 0.00065344 | 0.059661934 |
| Ern2 | 5.321418571 | 0.00065507 | 0.059661934 |
| Eps8l3 | 4.8750311 | 0.00067007 | 0.059661934 |
| Agr2 | 6.408416842 | 0.0006791 | 0.06000089 |
| Atp10b | 5.659373284 | 0.00071596 | 0.061875404 |
| Car8 | 4.740726037 | 0.00071608 | 0.061875404 |
| Myo6 | 1.712188398 | 0.00071648 | 0.061875404 |
| Flnb | 2.660341623 | 0.00073818 | 0.062444724 |
| Mettl7b | 6.347805578 | 0.00074071 | 0.062444724 |
| Cyp2c55 | 5.443939366 | 0.00074544 | 0.062444724 |
| Dgkh | 5.038798466 | 0.00077678 | 0.062444724 |
| Nr4a2 | 1.306791625 | 0.00078721 | 0.062444724 |
| Pld1 | 2.832337028 | 0.00079376 | 0.062444724 |
| Eps8l2 | 5.512005836 | 0.00080085 | 0.062444724 |
| Ptprk | 4.728602323 | 0.00080251 | 0.062444724 |
| Clca3b | 7.043129697 | 0.00080564 | 0.062444724 |
| Slc6a14 | 4.829980234 | 0.00080634 | 0.062444724 |
| Clec2h | 4.74853059 | 0.00080858 | 0.062444724 |
| Cpd | 1.301279103 | 0.0008119 | 0.062444724 |
| Fam160a1 | 4.887792221 | 0.00081452 | 0.062444724 |
| Plxnb2 | 2.166078694 | 0.00081549 | 0.062444724 |
| C77080 | 1.504482163 | 0.00082173 | 0.062505849 |
| Hist1h4n | -1.127765982 | 0.00082885 | 0.06263281 |
| Ckmt1 | 5.753771739 | 0.00083481 | 0.062670723 |
| Fosl2 | 1.37412865 | 0.00084379 | 0.062933564 |
| Rab15 | 4.69779248 | 0.00085577 | 0.063415537 |
| Lad1 | 4.947057201 | 0.00086141 | 0.063423703 |
| Cldn7 | 3.952239136 | 0.0008931 | 0.063688945 |
| Baiap2l1 | 2.535537987 | 0.00090239 | 0.063688945 |

| | | | |
|---|---|---|---|
| Atp12a | 4.363222907 | 0.00091658 | 0.063688945 |
| Dsc2 | 3.972988205 | 0.00091926 | 0.063688945 |
| Heph | 4.653303415 | 0.00092003 | 0.063688945 |
| Casz1 | 1.7575531 | 0.00092688 | 0.063688945 |
| Rimbp3 | 1.50012937 | 0.00092689 | 0.063688945 |
| Car4 | 4.464414874 | 0.00092899 | 0.063688945 |
| Klf6 | 1.029790859 | 0.00093959 | 0.063688945 |
| Vipr1 | 4.445865344 | 0.00094117 | 0.063688945 |
| Cftr | 5.503026857 | 0.00095466 | 0.063688945 |
| Vdr | 1.677097035 | 0.00095688 | 0.063688945 |
| Cdkn1a | 2.076493684 | 0.00096003 | 0.063688945 |
| Fut2 | 6.065673174 | 0.00096134 | 0.063688945 |
| St14 | 4.713933286 | 0.00096573 | 0.063688945 |
| Krt7 | 4.521284318 | 0.00096732 | 0.063688945 |
| Hpgd | 6.014471563 | 0.00096762 | 0.063688945 |
| Trpm6 | 4.480473718 | 0.00097391 | 0.063688945 |
| 5330417C22Rik | 4.088424382 | 0.00098861 | 0.063793119 |
| Ush1c | 5.625571623 | 0.0010154 | 0.065095092 |
| Rbm47 | 4.188386777 | 0.00102012 | 0.065095092 |
| Shtn1 | 4.404043904 | 0.00103248 | 0.065519959 |
| Ccl6 | 4.621312353 | 0.00106094 | 0.066955805 |
| Ehf | 5.072065409 | 0.00106898 | 0.067094404 |
| Aqp8 | 4.751071722 | 0.00109089 | 0.067365295 |
| 1810065E05Rik | 5.337820174 | 0.00110186 | 0.067379773 |
| Parva | 5.470724138 | 0.00110822 | 0.067379773 |
| Sema5a | 6.203923637 | 0.00110872 | 0.067379773 |
| Slfn4 | 5.262387034 | 0.00111566 | 0.067444673 |
| Pitx1 | 4.70224493 | 0.00113356 | 0.068096613 |
| Hao2 | 5.102225469 | 0.00115596 | 0.068096613 |
| Olfr402 | 4.299646872 | 0.00115832 | 0.068096613 |
| Appl2 | 2.940919858 | 0.00116365 | 0.068096613 |
| F11r | 4.493644566 | 0.00117772 | 0.068096613 |
| Pls1 | 4.184754783 | 0.00118006 | 0.068096613 |
| Mllt4 | 1.527399428 | 0.00118051 | 0.068096613 |
| Trpm4 | 4.178563647 | 0.00118365 | 0.068096613 |
| Tgm3 | 4.954284522 | 0.00118573 | 0.068096613 |
| Anxa3 | 4.666694048 | 0.00120109 | 0.068127226 |
| Entpd8 | 5.020951255 | 0.00120597 | 0.068127226 |
| Bcar1 | 4.382908399 | 0.00121524 | 0.068127226 |
| Rin2 | 3.217188736 | 0.00121592 | 0.068127226 |
| Tmem30b | 4.793247042 | 0.00124226 | 0.069264912 |
| 1810011O10Rik | 4.544691208 | 0.00125595 | 0.069690119 |
| Dock1 | 3.141566663 | 0.00128018 | 0.070625802 |

| | | | |
|---|---|---|---|
| Bcas1 | 4.228938946 | 0.00128511 | 0.070625802 |
| Hsd17b2 | 5.472665692 | 0.00129497 | 0.070828671 |
| Igsf5 | 5.463776104 | 0.00130682 | 0.071138276 |
| L1cam | 3.794461481 | 0.00132054 | 0.071545949 |
| Trib1 | 2.510223374 | 0.00133912 | 0.072211898 |
| B3gnt7 | 1.999814732 | 0.00134724 | 0.072256131 |
| Nlrp6 | 3.993314413 | 0.00135252 | 0.072256131 |
| Cdc42bpb | 1.277779205 | 0.00139229 | 0.073705296 |
| Cldn3 | 4.323617211 | 0.00139248 | 0.073705296 |
| Slc5a8 | 4.410686737 | 0.00140914 | 0.074244951 |
| Actc1 | 4.408117542 | 0.00143872 | 0.075457253 |
| Slc44a4 | 4.963863216 | 0.00149823 | 0.077593226 |
| Gprc5a | 4.964869731 | 0.00149944 | 0.077593226 |
| Zan | 2.511051249 | 0.00150421 | 0.077593226 |
| Dpep1 | 4.2107514 | 0.00150647 | 0.077593226 |
| Pde9a | 4.410668936 | 0.00151417 | 0.077641586 |
| Cx3cl1 | 4.332225989 | 0.00153512 | 0.07777137 |
| Ppp1r1b | 6.267564801 | 0.00153601 | 0.07777137 |
| Tmc5 | 4.217447779 | 0.00154688 | 0.07777137 |
| Etl4 | 4.199456679 | 0.00157862 | 0.078493484 |
| Dsg2 | 6.717560832 | 0.00158854 | 0.078550545 |
| Spdef | 4.363183429 | 0.00159344 | 0.078550545 |
| Ildr1 | 5.919743803 | 0.00161213 | 0.078980396 |
| Gcnt3 | 5.915448135 | 0.00161889 | 0.078980396 |
| Retnlb | 6.758876965 | 0.00162825 | 0.078980396 |
| Neo1 | 5.783887137 | 0.00163187 | 0.078980396 |
| Ptprd | 5.649905856 | 0.00163715 | 0.078980396 |
| Fam135a | 4.391558738 | 0.00164342 | 0.078980396 |
| Prr15l | 3.954356842 | 0.0016539 | 0.079152987 |
| Creb3l1 | 6.241843375 | 0.00166945 | 0.079565531 |
| Src | 3.825543179 | 0.00167786 | 0.079635932 |
| 2200002D01Rik | 2.649992046 | 0.00168842 | 0.079807186 |
| F2rl1 | 4.700393303 | 0.00172531 | 0.080416522 |
| Ugt2b34 | 6.189220286 | 0.00172756 | 0.080416522 |
| Tppp | 3.552801867 | 0.00172822 | 0.080416522 |
| Adh1 | 6.289299289 | 0.00172931 | 0.080416522 |
| Btnl4 | 4.191557298 | 0.00178527 | 0.082684119 |
| Tuba3b | 2.711715064 | 0.00179629 | 0.082860338 |
| Spry4 | -1.562864209 | 0.00180419 | 0.082891698 |
| Acta1 | 2.891951802 | 0.00181141 | 0.082891915 |
| Itgb4 | 2.116819485 | 0.0018213 | 0.083013809 |
| Baiap2l2 | 4.155954754 | 0.00183484 | 0.083300237 |
| Ms4a8a | 6.095543024 | 0.001847 | 0.083522197 |

| | | | |
|---|---|---|---|
| Slc9a2 | 3.982495364 | 0.00186258 | 0.08367682 |
| B3galt5 | 5.255270325 | 0.00186499 | 0.08367682 |
| Thbs1 | 3.884843103 | 0.0018767 | 0.083874696 |
| Syne2 | 3.117184521 | 0.00193286 | 0.085723957 |
| Socs3 | 2.388988238 | 0.00193906 | 0.085723957 |
| B3gnt3 | 4.443816813 | 0.00194047 | 0.085723957 |
| Id1 | 4.205155023 | 0.00195474 | 0.086023558 |
| Hoxb13 | 5.354400609 | 0.00196722 | 0.086242326 |
| Cttnbp2nl | 3.92735121 | 0.00197688 | 0.08633644 |
| Lmo7 | 6.512690221 | 0.00202539 | 0.088119987 |
| Slc2a1 | 2.522554222 | 0.00205402 | 0.088697166 |
| Pmp22 | 5.767723736 | 0.00205766 | 0.088697166 |
| Ttn | 1.619961595 | 0.00206183 | 0.088697166 |
| Mlph | 4.028573972 | 0.00207899 | 0.088802107 |
| Rnase4 | 5.346581096 | 0.00207973 | 0.088802107 |
| Foxa1 | 3.996601964 | 0.00211427 | 0.089811728 |
| Cdh17 | 6.477707096 | 0.00212446 | 0.089811728 |
| Mgat4a | 0.940055224 | 0.00213497 | 0.089811728 |
| Misp | 5.089254487 | 0.0021406 | 0.089811728 |
| 6430548M08Rik | 4.763840426 | 0.00214834 | 0.089811728 |
| Ccnd1 | -1.328173634 | 0.00215029 | 0.089811728 |
| Cers6 | 3.507041666 | 0.00216103 | 0.089814448 |
| Hist2h4 | -0.962807991 | 0.00216599 | 0.089814448 |
| Hgfac | 2.847905941 | 0.00220283 | 0.090915794 |
| Plch2 | 3.138559763 | 0.00220839 | 0.090915794 |
| Cdhr5 | 5.352651043 | 0.00225588 | 0.092462911 |
| Tns4 | 5.228227429 | 0.00226207 | 0.092462911 |
| Il4ra | 1.485032241 | 0.00227909 | 0.092828561 |
| Papss2 | 2.142115958 | 0.00229245 | 0.093042713 |
| Kcnq1 | 5.535034956 | 0.00232002 | 0.093680065 |
| Fam83e | 2.143490253 | 0.00233975 | 0.093680065 |
| Ano1 | 3.252736512 | 0.00234866 | 0.093680065 |
| Sdc4 | 3.362009193 | 0.00234893 | 0.093680065 |
| Thbs4 | 4.293122845 | 0.00236158 | 0.093858421 |
| Dst | 1.988996817 | 0.00237521 | 0.094074786 |
| Nr5a2 | 4.866460502 | 0.0023893 | 0.09430768 |
| Slc26a3 | 5.093261279 | 0.00243709 | 0.095648102 |
| Galnt3 | 4.384674377 | 0.00243992 | 0.095648102 |
| Cmtm4 | 1.774108484 | 0.00249064 | 0.097304226 |
| Sh3bgrl2 | 3.270899628 | 0.00251183 | 0.097799603 |
| Spint1 | 3.986477052 | 0.00252579 | 0.098011017 |
| Hspa1l | 1.96644249 | 0.00253442 | 0.098014604 |
| Cds1 | 4.143409544 | 0.00256582 | 0.09860838 |

| | | | |
|---|---|---|---|
| Sptan1 | 2.399237354 | 0.00256694 | 0.09860838 |
| Itga3 | 2.700680684 | 0.00260982 | 0.098968766 |
| Pde3a | 4.341584788 | 0.00261456 | 0.098968766 |
| Fer1l6 | 4.836848398 | 0.00262045 | 0.098968766 |
| Aqp4 | 6.670541083 | 0.0026213 | 0.098968766 |
| Il21 | 3.593460151 | 0.00262506 | 0.098968766 |
| Kank1 | 4.178315906 | 0.00263426 | 0.098968766 |
| Slc9a3 | 4.886578893 | 0.00264487 | 0.098968766 |
| Tmcc3 | 2.61503398 | 0.00264526 | 0.098968766 |
| Mapk13 | 5.168807033 | 0.00266026 | 0.099206979 |
| Prom1 | 5.132009716 | 0.00270143 | 0.100239843 |
| Ctla2a | 2.200746347 | 0.00270541 | 0.100239843 |
| Robo4 | 4.533137438 | 0.00272133 | 0.100505396 |
| Efnb2 | 3.211402023 | 0.00273378 | 0.100641581 |
| Atp9a | 3.423492684 | 0.00274863 | 0.100814236 |
| Sult1a1 | 1.972173932 | 0.00275602 | 0.100814236 |
| Maoa | 1.304252499 | 0.0027994 | 0.10172017 |
| Lrp4 | 4.657781452 | 0.00280703 | 0.10172017 |
| Arhgef16 | 3.728028775 | 0.00280736 | 0.10172017 |
| Ppp1r3d | 3.677605326 | 0.0028278 | 0.102033729 |
| Fabp2 | 5.69913694 | 0.00283928 | 0.102033729 |
| Gucy2c | 4.82243051 | 0.00284266 | 0.102033729 |
| Cdx1 | 4.756179441 | 0.00291712 | 0.104056162 |
| Fosb | 0.792938122 | 0.00294713 | 0.104477526 |
| Epas1 | 3.296332092 | 0.00295645 | 0.104485613 |
| Apc2 | 1.97854957 | 0.00296882 | 0.104600785 |
| Egr3 | 1.862397983 | 0.00299067 | 0.104606545 |
| Errfi1 | 3.901162644 | 0.0029963 | 0.104606545 |
| Hbegf | 2.72105618 | 0.00304393 | 0.105947186 |
| Isx | 3.627178798 | 0.00312648 | 0.108164735 |
| Tmem45b | 3.344402313 | 0.00316557 | 0.109188446 |
| Aldob | 4.401350295 | 0.00320622 | 0.11025949 |
| Yes1 | 3.94324146 | 0.00323719 | 0.110581444 |
| Lrig3 | 3.219551577 | 0.00324447 | 0.110581444 |
| Tcf7l2 | 2.16456386 | 0.00327046 | 0.111137494 |
| Sult1d1 | 5.208371644 | 0.0032971 | 0.11120658 |
| Plekhg3 | 2.826605702 | 0.00329938 | 0.11120658 |
| Shroom2 | 3.702073003 | 0.00330154 | 0.11120658 |
| Sorl1 | 2.330292898 | 0.00337707 | 0.112763359 |
| Frk | 5.398347516 | 0.00337884 | 0.112763359 |
| Pcsk5 | 5.461095695 | 0.0033882 | 0.112763359 |
| Zg16 | 1.264989969 | 0.00340321 | 0.112763359 |
| Lamb3 | 5.374012048 | 0.00340666 | 0.112763359 |

| | | | |
|---|---|---|---|
| Tm4sf20 | 3.612695232 | 0.00342543 | 0.113058803 |
| Cpeb2 | 1.545578841 | 0.00350755 | 0.115002391 |
| Slc26a1 | -1.118330222 | 0.00350829 | 0.115002391 |
| Sytl2 | 1.877754996 | 0.00351435 | 0.115002391 |
| Arhgef28 | 3.097330812 | 0.00353652 | 0.115156084 |
| Tpd52 | 1.238106372 | 0.00354913 | 0.115156084 |
| Chmp4c | 3.585325341 | 0.0035768 | 0.115727016 |
| Map3k9 | 3.767708517 | 0.00360967 | 0.116462611 |
| Abr | 1.146253703 | 0.00362942 | 0.116771626 |
| Fzd7 | -0.71485864 | 0.00366877 | 0.117708052 |
| Dgka | 1.925183898 | 0.00370046 | 0.118394022 |
| 2010300C02Rik | 3.937471621 | 0.00373827 | 0.119059432 |
| Tnks1bp1 | 1.66898444 | 0.00374199 | 0.119059432 |
| Nr3c2 | 2.463129108 | 0.00377985 | 0.119932001 |
| Gfod1 | 1.915750276 | 0.00380661 | 0.12022812 |
| Ptgfrn | 3.224170594 | 0.00381012 | 0.12022812 |
| Tubb4a | 3.280628556 | 0.00385371 | 0.120411012 |
| Abat | 2.557041872 | 0.00385584 | 0.120411012 |
| Atp2c2 | 3.483560265 | 0.00385785 | 0.120411012 |
| Sox9 | 3.294204559 | 0.00389027 | 0.121093718 |
| Galnt12 | 4.100220501 | 0.00392001 | 0.121689891 |
| Pim1 | 0.799248955 | 0.00393493 | 0.121823769 |
| Tbc1d2 | 1.796404968 | 0.00395483 | 0.122110727 |
| Cnnm4 | 1.318845788 | 0.00399935 | 0.12315415 |
| Pcnx | 0.75489889 | 0.00402982 | 0.12343083 |
| Uaca | 3.866220171 | 0.00405313 | 0.12371886 |
| Apob | 5.33234319 | 0.0040616 | 0.12371886 |
| Aim1 | 1.756573881 | 0.00410335 | 0.124029143 |
| Egr1 | 0.668177307 | 0.0041343 | 0.124636545 |
| Col5a1 | -0.705651174 | 0.004178 | 0.124998542 |
| Gab2 | 1.066006046 | 0.00417895 | 0.124998542 |
| Hist1h2ae | -0.844448909 | 0.00422589 | 0.125965081 |
| Prss32 | 5.934552964 | 0.0042332 | 0.125965081 |
| Cdcp1 | 2.460578786 | 0.00427656 | 0.126642616 |
| Fam83h | 2.419736033 | 0.00429218 | 0.126642616 |
| Farp1 | 1.852429328 | 0.0042923 | 0.126642616 |
| Cdx2 | 4.013951775 | 0.00431018 | 0.126642616 |
| Tiparp | 1.316434586 | 0.00433629 | 0.126734509 |
| Eps8 | 3.719745566 | 0.00442785 | 0.129081989 |
| Fam3b | 3.25820342 | 0.00450766 | 0.13074503 |
| Cyp2c65 | 4.343080379 | 0.00456933 | 0.131867654 |
| Ano9 | 3.665397487 | 0.00463541 | 0.133439441 |
| Pcdh1 | 3.560907977 | 0.00469944 | 0.134944444 |

| | | | |
|---|---|---|---|
| Fmn1 | 3.460073858 | 0.00473502 | 0.135368919 |
| Sel1l3 | 2.867999914 | 0.00473779 | 0.135368919 |
| Aaas | -0.653909326 | 0.00477498 | 0.136011634 |
| Zfp467 | 3.164346344 | 0.00478397 | 0.136011634 |
| Raph1 | 2.13775386 | 0.00480353 | 0.136230426 |
| Rorc | 2.864222005 | 0.00493004 | 0.139414335 |
| Hmgcs1 | 0.753242105 | 0.00494007 | 0.139414335 |
| Sash1 | 2.276783475 | 0.00497098 | 0.139942858 |
| Atp2a3 | 2.126866603 | 0.00500042 | 0.140427412 |
| Plce1 | 3.43616176 | 0.0050175 | 0.140429235 |
| Mst1r | 5.034581238 | 0.00513328 | 0.142903264 |
| Mecom | 3.264649426 | 0.00513835 | 0.142903264 |
| Fmnl3 | 1.496144723 | 0.0051873 | 0.143916108 |
| Cd177 | 4.906910613 | 0.00531823 | 0.146487119 |
| Plekha7 | 1.53454906 | 0.00542922 | 0.149186523 |
| Itga2 | 2.303968108 | 0.00548291 | 0.149679483 |
| Gpsm1 | 1.622752962 | 0.00550787 | 0.149679483 |
| Bspry | 2.867533312 | 0.00551869 | 0.149679483 |
| Hist1h1e | -0.664831828 | 0.00552374 | 0.149679483 |
| Gipc2 | 5.048441336 | 0.00552534 | 0.149679483 |
| Hist1h2ab | -0.888541811 | 0.00557314 | 0.150618919 |
| Col23a1 | 0.87105337 | 0.00559262 | 0.150790551 |
| H2-Bl | -0.621646334 | 0.00565781 | 0.151684621 |
| Rab25 | 3.228175607 | 0.0056786 | 0.151684621 |
| Hist1h2bj | -0.885384463 | 0.00570605 | 0.152064353 |
| Hist1h2bl | -0.750097946 | 0.00581494 | 0.154333525 |
| Mboat1 | 2.519733346 | 0.00581808 | 0.154333525 |
| Hist1h2bp | -0.869770205 | 0.00584361 | 0.154653811 |
| Clic5 | 4.224140336 | 0.00586482 | 0.154858145 |
| Kcne3 | 2.709332622 | 0.00588061 | 0.154919081 |
| Thrb | 3.067604731 | 0.00590946 | 0.155322692 |
| Setbp1 | 2.479042228 | 0.00601045 | 0.157616524 |
| Degs2 | 4.060900554 | 0.00607171 | 0.158315631 |
| Acsf2 | 0.997719811 | 0.00607846 | 0.158315631 |
| Arhgef10l | 3.196203858 | 0.00612604 | 0.158987567 |
| Fam114a1 | 2.793561936 | 0.0061368 | 0.158987567 |
| Arfgef3 | 3.521671535 | 0.00614578 | 0.158987567 |
| As3mt | 2.069495601 | 0.00624272 | 0.161132398 |
| Arhgef40 | 1.079561697 | 0.00642114 | 0.164627591 |
| Coro2a | 2.621650043 | 0.0064437 | 0.164838088 |
| Prkcq | 1.374821808 | 0.00646274 | 0.164957919 |
| Slc39a4 | 2.531459455 | 0.0065195 | 0.165649174 |
| Ctsf | 2.439710805 | 0.00652205 | 0.165649174 |

| | | | |
|---|---|---|---|
| Rap1gap2 | 1.899910556 | 0.00653309 | 0.165649174 |
| Ddit4 | 1.124916342 | 0.00657708 | 0.166224277 |
| Dip2a | 1.379139503 | 0.00658472 | 0.166224277 |
| Cldn2 | 5.158646154 | 0.00665382 | 0.167600338 |
| 5-Sep | 3.454092834 | 0.0066902 | 0.168147962 |
| Cplx2 | 1.259809162 | 0.00678556 | 0.169801693 |
| Hist1h2ac | -0.783958044 | 0.00682197 | 0.170170995 |
| Myof | 4.348701391 | 0.00682995 | 0.170170995 |
| Btg2 | 0.713099901 | 0.00687129 | 0.170830272 |
| Rasa4 | 2.251676647 | 0.00692801 | 0.171868456 |
| Pf4 | 1.943374379 | 0.00699091 | 0.17305516 |
| Inadl | 2.358881399 | 0.00702078 | 0.173246162 |
| Epb41l1 | 2.769138806 | 0.00704322 | 0.173246162 |
| Rassf6 | 2.936177125 | 0.00704388 | 0.173246162 |
| Hspa1b | 1.686782011 | 0.00709293 | 0.174079931 |
| Perp | 2.962048789 | 0.007228 | 0.176557557 |
| Pglyrp1 | 2.788156479 | 0.00724 | 0.176557557 |
| Cldn15 | 3.200982883 | 0.00726185 | 0.17659237 |
| Atrnl1 | 1.140557311 | 0.00732323 | 0.177297822 |
| Ifi27l2b | 5.220326078 | 0.00737835 | 0.177513107 |
| Lgr4 | 1.487673044 | 0.00738334 | 0.177513107 |
| Gpr132 | 1.294766274 | 0.00745299 | 0.178343899 |
| Klf1 | -0.569824496 | 0.00750192 | 0.179038556 |
| Hist1h3d | -0.795828869 | 0.0075132 | 0.179038556 |
| Themis3 | 4.827724161 | 0.00755298 | 0.179614011 |
| Pappa | -0.603560904 | 0.0075984 | 0.180320702 |
| Nupr1 | 3.342102571 | 0.00772177 | 0.182494395 |
| Srxn1 | 1.250537734 | 0.00784331 | 0.184606962 |
| Ehd3 | 1.980130567 | 0.00795502 | 0.186853394 |
| Serpinb6a | 1.237951462 | 0.0080432 | 0.188539109 |
| Gas6 | 3.436404845 | 0.00811556 | 0.189847936 |
| Hist1h1b | -0.728703946 | 0.00818444 | 0.191070062 |
| Sptbn1 | 1.617982771 | 0.00831338 | 0.192893721 |
| Slc30a4 | 3.175493848 | 0.00832872 | 0.192893721 |
| Cactin | -0.8649571 | 0.00832973 | 0.192893721 |
| Bmpr1a | 4.857455507 | 0.00835507 | 0.193091309 |
| Mylk | 2.867876463 | 0.0084 | 0.193739689 |
| Plk2 | 2.165695252 | 0.00842041 | 0.193821236 |
| Serinc2 | 4.759200706 | 0.00852626 | 0.195865146 |
| Dusp5 | 1.107267763 | 0.00854883 | 0.19599169 |
| Acsm3 | 2.096907473 | 0.00869315 | 0.198903341 |
| Lpcat2 | 2.56018138 | 0.00882359 | 0.201486552 |
| Slc39a5 | 2.912915408 | 0.00889497 | 0.202713559 |

| | | | |
|---|---|---|---|
| Vldlr | 5.590806226 | 0.00892213 | 0.202917854 |
| Anxa2 | 1.316005957 | 0.00894232 | 0.202917854 |
| Clca4a | 3.521454558 | 0.00895694 | 0.202917854 |
| Ccnf | -0.634726275 | 0.00899418 | 0.20336049 |
| Slc5a1 | 2.786456954 | 0.00901984 | 0.203539977 |
| Npas2 | 3.477449178 | 0.00906023 | 0.203725933 |
| Neb | 2.140838534 | 0.00906355 | 0.203725933 |
| Irs2 | 0.593668477 | 0.00918297 | 0.205868015 |
| Slc35d3 | 3.500927724 | 0.0091947 | 0.205868015 |
| Aim1l | 3.481853998 | 0.00925902 | 0.206658922 |
| Tead3 | 1.151639925 | 0.00926601 | 0.206658922 |
| Smad7 | 0.831028212 | 0.0093096 | 0.207228795 |
| Ddah1 | 2.844523272 | 0.00941349 | 0.208732422 |
| Cdc6 | -0.551107805 | 0.00945465 | 0.20921207 |
| Plekhb2 | 0.801750584 | 0.0094797 | 0.20921207 |
| Pawr | 2.569947934 | 0.00948977 | 0.20921207 |
| Zfp36 | 0.925171075 | 0.00959117 | 0.210482994 |
| Mmp15 | 2.906570728 | 0.00960239 | 0.210482994 |
| Fam234a | 0.923853381 | 0.00972946 | 0.212770155 |
| Mical2 | 2.796251005 | 0.00974378 | 0.212770155 |
| Calml4 | 2.09138696 | 0.00978106 | 0.212861746 |
| 1700027J19Rik | 3.561162275 | 0.00980415 | 0.212861746 |
| Cyr61 | 2.335046902 | 0.00980703 | 0.212861746 |
| Rap1b | 0.556381975 | 0.00982211 | 0.212861746 |
| Cd38 | 3.144275933 | 0.00986361 | 0.213358571 |
| Ephb2 | 4.207202876 | 0.00992021 | 0.214179491 |
| Tuba3a | 3.481594046 | 0.01000654 | 0.215638047 |
| Itih5 | 2.988182067 | 0.01014324 | 0.217360631 |
| Gramd3 | 3.255911557 | 0.01026529 | 0.219165182 |
| Hist1h2be | -0.710897543 | 0.01026562 | 0.219165182 |
| Plek | 2.215771078 | 0.01033374 | 0.219723286 |
| Hist1h1c | -0.653282891 | 0.01036142 | 0.219723286 |
| Sema4a | 2.347939401 | 0.01036828 | 0.219723286 |
| Plxnd1 | 1.790642264 | 0.01042323 | 0.220265054 |
| Mef2c | 1.257807322 | 0.0104322 | 0.220265054 |
| Fhl1 | 3.510284125 | 0.01048776 | 0.221031951 |
| Hist1h4c | -0.794636482 | 0.01057622 | 0.222422469 |
| Zcchc14 | 1.861114634 | 0.01059247 | 0.222422469 |
| Tcf7 | 1.33378654 | 0.01065589 | 0.223268857 |
| Fnbp1l | 1.200180143 | 0.01067165 | 0.223268857 |
| Ugp2 | 0.940839968 | 0.01074758 | 0.224448558 |
| Spryd4 | -0.961400141 | 0.01079688 | 0.225068876 |
| Smpd3 | 1.590107731 | 0.01082483 | 0.225161755 |

| | | | |
|---|---|---|---|
| Ppp1r9b | -0.623193335 | 0.01084054 | 0.225161755 |
| Dennd1c | 1.880615522 | 0.0109092 | 0.2255635 |
| Sdc1 | 1.704786545 | 0.01091347 | 0.2255635 |
| Prkca | 1.498221237 | 0.0109188 | 0.2255635 |
| Hist2h2bb | -0.865496639 | 0.01109571 | 0.228105708 |
| Arhgef25 | -1.169002687 | 0.01110144 | 0.228105708 |
| Hsd11b2 | 2.377387018 | 0.01114775 | 0.228648227 |
| Cenpf | -0.522594223 | 0.0112264 | 0.229851015 |
| Plekho1 | 1.329784908 | 0.01130547 | 0.23072381 |
| Fgd4 | 2.518912079 | 0.01143566 | 0.232623407 |
| Tpm4 | 1.450110688 | 0.01145604 | 0.232623407 |
| Med12l | 2.013033768 | 0.01146307 | 0.232623407 |
| Plaur | 2.465473656 | 0.01164832 | 0.235550292 |
| Pard3b | 1.098119554 | 0.01171369 | 0.235968512 |
| Camk2n1 | 2.634057527 | 0.01175777 | 0.236100845 |
| Eml4 | 0.836188711 | 0.01185717 | 0.236854605 |
| F2r | 0.98510554 | 0.01195749 | 0.237618887 |
| Fanca | -0.552182846 | 0.01198983 | 0.237850059 |
| Sphk1 | -1.072431919 | 0.01204258 | 0.238484518 |
| Sorbs3 | 2.677969783 | 0.01208902 | 0.238992172 |
| Wdr60 | -0.727217139 | 0.01221318 | 0.241031853 |
| Dok4 | 2.029295826 | 0.01229626 | 0.242255328 |
| Zfp637 | -0.792253437 | 0.01233051 | 0.242447907 |
| Lat | 1.133297727 | 0.01234825 | 0.242447907 |
| Pde4b | 0.864622415 | 0.012426 | 0.242749512 |
| Snx33 | 1.17660774 | 0.01242702 | 0.242749512 |
| Inppl1 | 1.160012176 | 0.0125193 | 0.243723114 |
| Tdrd7 | 0.787937838 | 0.01261814 | 0.245231744 |
| Sgpp2 | 1.476526079 | 0.01267664 | 0.245564951 |
| Adgra3 | 1.162842936 | 0.01269071 | 0.245564951 |
| Rab27b | 1.668494487 | 0.01269942 | 0.245564951 |
| Cad | -0.525212655 | 0.0127231 | 0.245609266 |
| Muc13 | 1.07988614 | 0.01280393 | 0.246404236 |
| Mxd1 | 0.679650491 | 0.01282648 | 0.246404236 |
| Slc23a2 | 0.811864773 | 0.01282864 | 0.246404236 |
| 4930539E08Rik | 3.342662401 | 0.01286682 | 0.24672502 |
| Rhpn2 | 3.893297804 | 0.01292545 | 0.246962274 |
| Tmc4 | 1.101159926 | 0.0129942 | 0.246962274 |
| Ola1 | -0.543879488 | 0.0130007 | 0.246962274 |
| Erbb3 | 1.111933423 | 0.01300803 | 0.246962274 |
| Myadm | 2.499162568 | 0.0130099 | 0.246962274 |
| Rap1gap | 2.046197562 | 0.01301246 | 0.246962274 |
| Spint2 | 0.609153171 | 0.0130297 | 0.246962274 |

| | | | |
|---|---|---|---|
| Slc7a6 | 1.287091072 | 0.01306762 | 0.24704374 |
| Etv5 | -0.927739892 | 0.01307702 | 0.24704374 |
| Arap2 | 0.892377545 | 0.01314115 | 0.247430231 |
| Nfatc2 | 2.148092005 | 0.01314444 | 0.247430231 |
| Ephb4 | 1.894243874 | 0.0131621 | 0.247430231 |
| Nt5e | 2.481175179 | 0.01318865 | 0.247524208 |
| Hist1h2ak | -0.680258314 | 0.01333078 | 0.249376741 |
| Vps13c | 0.802423078 | 0.01335627 | 0.249447299 |
| 8-Sep | -0.657115609 | 0.01354299 | 0.252523961 |
| Chn2 | 2.40905723 | 0.01363395 | 0.253397338 |
| Mns1 | -1.035292067 | 0.01367802 | 0.253805764 |
| Magi1 | 1.624532179 | 0.0137502 | 0.254733476 |
| Cgref1 | 2.532788907 | 0.01380953 | 0.255420727 |
| Smim24 | 1.470351984 | 0.01388256 | 0.255657789 |
| Rtkn | 4.542385324 | 0.01391977 | 0.255657789 |
| Hist3h2a | -0.85398313 | 0.01393632 | 0.255657789 |
| Tubb3 | 1.724468584 | 0.01394922 | 0.255657789 |
| Tmem44 | 2.318237725 | 0.0139559 | 0.255657789 |
| Hist2h2ac | -0.774719244 | 0.01401704 | 0.256368904 |
| Atp8b2 | 1.467370259 | 0.01410796 | 0.257621665 |
| Atf3 | 1.33789723 | 0.01417131 | 0.258019479 |
| Trpc6 | 2.642804145 | 0.01417467 | 0.258019479 |
| Csrp2 | 2.22735026 | 0.01426108 | 0.258988689 |
| Tgoln1 | 0.52070495 | 0.01427301 | 0.258988689 |
| Tjp3 | 1.02919495 | 0.0143777 | 0.260276735 |
| Nhsl1 | 1.989582868 | 0.01438932 | 0.260276735 |
| Pkm | 1.780257759 | 0.01442069 | 0.260434017 |
| Ano7 | 2.11918434 | 0.01455814 | 0.262503589 |
| Spdl1 | -0.570149868 | 0.01461221 | 0.263065569 |
| Sun2 | 0.554826992 | 0.01465426 | 0.263409788 |
| Ugdh | 0.706347466 | 0.01476763 | 0.265032879 |
| Arl4c | 1.145938441 | 0.01486719 | 0.265350675 |
| Hist1h2bh | -0.786201273 | 0.01489408 | 0.265350675 |
| Tjp2 | 1.057344979 | 0.0149359 | 0.265350675 |
| Bard1 | -0.750178447 | 0.01493629 | 0.265350675 |
| Ano10 | 2.156911307 | 0.01494812 | 0.265350675 |
| Mcm10 | -0.608476191 | 0.01503226 | 0.265350675 |
| Acta2 | 2.433529086 | 0.01503915 | 0.265350675 |
| Hist1h4a | -0.782931751 | 0.01505379 | 0.265350675 |
| Rel | 1.205857687 | 0.01506756 | 0.265350675 |
| Pdxdc1 | 0.525294652 | 0.01511176 | 0.265350675 |
| Ldlr | 0.517882044 | 0.01512469 | 0.265350675 |
| Nr4a3 | 2.226792274 | 0.01513187 | 0.265350675 |

| | | | |
|---|---|---|---|
| Arhgap32 | 2.482757148 | 0.01518274 | 0.265836762 |
| Xpo7 | 0.834236575 | 0.01531639 | 0.26748292 |
| Ctss | 2.007964537 | 0.01532333 | 0.26748292 |
| Rora | 2.529852631 | 0.01535755 | 0.267673423 |
| Mdfic | 2.235628515 | 0.01538585 | 0.267760385 |
| Ablim1 | 1.979994902 | 0.01542288 | 0.26799881 |
| Lrp12 | 0.970554703 | 0.01546612 | 0.268231157 |
| Nuf2 | -0.664278579 | 0.01549976 | 0.268231157 |
| Tbc1d4 | 1.30593589 | 0.01553516 | 0.268231157 |
| Igf2bp2 | 1.719599103 | 0.0155731 | 0.268231157 |
| Tbc1d10b | -0.487436146 | 0.01560363 | 0.268231157 |
| Dync2h1 | 0.649421467 | 0.01564222 | 0.268231157 |
| Ptbp2 | 0.883287746 | 0.01567058 | 0.268231157 |
| Cxadr | 1.096963121 | 0.01567734 | 0.268231157 |
| Hist1h2bb | -0.835916347 | 0.01569313 | 0.268231157 |
| Fam102a | 2.130883379 | 0.01574894 | 0.268368209 |
| Dnajc1 | 0.564409212 | 0.01575461 | 0.268368209 |
| Etv3 | 0.620760488 | 0.01579711 | 0.268368209 |
| Hist1h4d | -0.64112965 | 0.01580712 | 0.268368209 |
| Inpp4a | 1.166532797 | 0.01581798 | 0.268368209 |
| Cuedc1 | 1.840367384 | 0.01594537 | 0.270130524 |
| Serpinb1a | 2.85936963 | 0.01601911 | 0.270754126 |
| Sestd1 | 1.60021866 | 0.01602932 | 0.270754126 |
| Kif20a | -0.641613522 | 0.01609761 | 0.271508211 |
| Lamb2 | 3.018599107 | 0.01635831 | 0.275500862 |
| Mansc1 | 2.591129562 | 0.0165832 | 0.278879369 |
| Smim1 | -0.7466576 | 0.0166719 | 0.279749895 |
| Ddr1 | 2.802187054 | 0.01668367 | 0.279749895 |
| Psd3 | 0.867110702 | 0.01677351 | 0.280819085 |
| Hist1h2bm | -0.692751861 | 0.01679634 | 0.280819085 |
| Bag2 | -0.636677367 | 0.01695057 | 0.281235478 |
| Capn5 | 0.773524647 | 0.01699449 | 0.281235478 |
| Zyx | 1.081040721 | 0.01703236 | 0.281235478 |
| Pycard | 0.721875524 | 0.01705108 | 0.281235478 |
| Plekha5 | 0.740577167 | 0.0170561 | 0.281235478 |
| Kbtbd11 | 1.963984431 | 0.01706645 | 0.281235478 |
| Cul4a | -0.469957518 | 0.0170898 | 0.281235478 |
| Plk3 | 0.881247481 | 0.01709 | 0.281235478 |
| B4galt6 | 2.574984823 | 0.01709607 | 0.281235478 |
| Dhx37 | -0.466282593 | 0.01711506 | 0.281235478 |
| Lrrc75a | 0.959629208 | 0.01721578 | 0.282486281 |
| Klhl12 | -0.511897659 | 0.01729056 | 0.282709579 |
| Blnk | 1.766475074 | 0.01731533 | 0.282709579 |

| | | | |
|---|---|---|---|
| Mettl21c | 2.24654338 | 0.01733018 | 0.282709579 |
| Bambi | 1.962792416 | 0.01736879 | 0.282709579 |
| Scarf1 | 2.839595837 | 0.01738782 | 0.282709579 |
| Zfp516 | 1.699681836 | 0.01738867 | 0.282709579 |
| Plxna2 | 1.571558222 | 0.01740168 | 0.282709579 |
| 2310033P09Rik | -0.675968527 | 0.01752626 | 0.284331349 |
| Fam110c | 1.749528014 | 0.01757952 | 0.284577549 |
| Tmem40 | 0.732108579 | 0.01761576 | 0.284577549 |
| Cish | 1.718983922 | 0.01765697 | 0.284842662 |
| Il17rc | 2.915804766 | 0.01773259 | 0.285661393 |
| Kirrel3 | 1.296597074 | 0.01776295 | 0.285749617 |
| Selp | 3.932925465 | 0.01785483 | 0.286665981 |
| Hexim2 | -1.750860267 | 0.01786983 | 0.286665981 |
| Aldh1l1 | 1.515651012 | 0.01798445 | 0.287893397 |
| Wls | 1.246516231 | 0.01799647 | 0.287893397 |
| Zfp948 | 1.469865647 | 0.01803355 | 0.288085313 |
| Epsti1 | 1.69785288 | 0.01807014 | 0.288268968 |
| Tie1 | 2.324841334 | 0.01810217 | 0.288379307 |
| Prkcz | 1.924709006 | 0.01837056 | 0.291046486 |
| Dmwd | -0.55317915 | 0.01837943 | 0.291046486 |
| Utp14b | 0.804408606 | 0.01838342 | 0.291046486 |
| Mctp2 | 2.205412205 | 0.01841788 | 0.291046486 |
| Acvr2a | 1.042514119 | 0.01842081 | 0.291046486 |
| F2rl2 | 0.917238311 | 0.01842163 | 0.291046486 |
| Hist1h4k | -0.788409326 | 0.01866617 | 0.293766115 |
| Hspb1 | 4.209510468 | 0.01868736 | 0.293766115 |
| Sytl1 | 1.928874301 | 0.01869112 | 0.293766115 |
| Runx1 | 0.607216043 | 0.018716 | 0.293766115 |
| Lsr | 1.566873883 | 0.01872164 | 0.293766115 |
| Polrmt | -0.486509243 | 0.01891568 | 0.296405853 |
| Osm | 1.438748986 | 0.01896792 | 0.296819493 |
| Eya1 | 1.620693978 | 0.01899923 | 0.29690493 |
| Arhgef3 | 1.370040735 | 0.01913343 | 0.298595812 |
| Pold1 | -0.460259941 | 0.0192231 | 0.299588239 |
| Mmrn1 | 2.989124137 | 0.01927217 | 0.299946028 |
| Rnasel | 0.679045823 | 0.01938268 | 0.301166891 |
| Map3k13 | 1.222471798 | 0.01940927 | 0.301166891 |
| Pcsk9 | 1.257713602 | 0.01942928 | 0.301166891 |
| Hist1h4j | -0.738758381 | 0.01956531 | 0.301815096 |
| Lrrc47 | -0.46268999 | 0.01959495 | 0.301815096 |
| B3gnt5 | 2.103738814 | 0.01960276 | 0.301815096 |
| Slc25a44 | -0.549657255 | 0.01963758 | 0.301815096 |
| Ankrd61 | 1.811261942 | 0.01965503 | 0.301815096 |

| | | | |
|---|---|---|---|
| H2afx | -0.577535212 | 0.01974856 | 0.302846349 |
| Sgk1 | 1.591894968 | 0.01981906 | 0.303522322 |
| Greb1l | 1.595404927 | 0.01994707 | 0.305075994 |
| Cpeb4 | 0.913572481 | 0.02011753 | 0.307273831 |
| Trp53inp2 | 0.979584034 | 0.02014935 | 0.307351238 |
| Gm7694 | 2.422417885 | 0.02020254 | 0.307650592 |
| Plcb3 | 0.723499157 | 0.02022255 | 0.307650592 |
| Scand1 | -0.530074378 | 0.02027746 | 0.308077897 |
| Slc26a2 | 0.719113329 | 0.02032579 | 0.308404325 |
| Sf3a2 | -0.463822548 | 0.02037792 | 0.308631585 |
| Hist1h4f | -0.77600008 | 0.02046109 | 0.30923167 |
| Sort1 | 0.758042255 | 0.02055241 | 0.310203671 |
| Coq9 | -0.509435561 | 0.02076859 | 0.312404672 |
| Fbxl22 | 1.176041585 | 0.02077983 | 0.312404672 |
| Ahr | 1.520761608 | 0.02094996 | 0.314550643 |
| Fry | 0.962033873 | 0.02097798 | 0.314560144 |
| Gmds | 0.730002507 | 0.02102187 | 0.31480722 |
| Ralgds | 1.333909952 | 0.02107518 | 0.315194702 |
| Clock | 0.712489784 | 0.0211216 | 0.315478098 |
| Vwf | 1.030201282 | 0.02116456 | 0.315709247 |
| Rbm10 | -0.439554107 | 0.02128731 | 0.317077344 |
| Lonrf3 | 2.535971609 | 0.02131148 | 0.317077344 |
| Ifrd2 | -0.575681017 | 0.02160881 | 0.32067026 |
| Tmem2 | 0.799185031 | 0.02170067 | 0.321453711 |
| Amigo2 | -0.627668828 | 0.02174052 | 0.321453711 |
| Lrrn3 | 1.921264286 | 0.02174556 | 0.321453711 |
| Espl1 | -0.610675186 | 0.02178741 | 0.321658301 |
| Xdh | 1.540946563 | 0.02192599 | 0.323288788 |
| Smagp | 1.697097162 | 0.02196671 | 0.323473813 |
| Parp12 | 0.827297415 | 0.02214555 | 0.325689793 |
| Slc6a15 | 2.272680299 | 0.0223328 | 0.328023663 |
| Gpr171 | 2.747683254 | 0.02240494 | 0.328134553 |
| Sh3pxd2a | 1.249427687 | 0.02242416 | 0.328134553 |
| Aplp2 | -0.594603758 | 0.02265684 | 0.329981518 |
| Chdh | 3.307066092 | 0.02266372 | 0.329981518 |
| F2rl3 | 1.437590844 | 0.02268792 | 0.329981518 |
| Steap3 | -0.450759665 | 0.02269593 | 0.329981518 |
| Nbeal1 | 0.602091817 | 0.02275485 | 0.330419907 |
| 2310036O22Rik | -0.57362077 | 0.02284611 | 0.330940368 |
| Gfpt2 | 2.137329558 | 0.02287324 | 0.330940368 |
| Itga2b | 1.539317036 | 0.02287712 | 0.330940368 |
| Atp10a | 1.305222583 | 0.02295768 | 0.331687989 |
| Pip4k2c | -0.504667288 | 0.02302558 | 0.332251025 |

| | | | |
|---|---|---|---|
| Slc27a1 | 1.442305453 | 0.02308329 | 0.332665758 |
| Hist2h2ab | -0.609231406 | 0.0231338 | 0.332769814 |
| Slfn5 | 1.890432943 | 0.02325072 | 0.333822181 |
| Cbs | 1.977275863 | 0.02336022 | 0.334975583 |
| Pik3c2a | 1.664683631 | 0.02342721 | 0.335517359 |
| Tubb6 | 1.313587284 | 0.02352987 | 0.336568 |
| Mapk3 | 0.562954498 | 0.02363405 | 0.337158017 |
| Hist2h3b | -0.711980773 | 0.02364219 | 0.337158017 |
| Cachd1 | 1.330059546 | 0.02365918 | 0.337158017 |
| Tmem176a | 3.130093575 | 0.02369213 | 0.337209186 |
| Pcdhgc3 | 1.868978178 | 0.02383573 | 0.337939209 |
| Ldlrad3 | 1.192455391 | 0.02386137 | 0.337939209 |
| Nedd4l | 0.72489571 | 0.02391943 | 0.337939209 |
| Cdt1 | -0.591950991 | 0.02393166 | 0.337939209 |
| Muc5b | 1.63488277 | 0.0239424 | 0.337939209 |
| Pbx1 | 1.606595471 | 0.02397727 | 0.337939209 |
| Irf6 | 1.641702991 | 0.02397825 | 0.337939209 |
| Knstrn | -0.538108716 | 0.0239788 | 0.337939209 |
| Nphp1 | -0.772375144 | 0.02408301 | 0.338535791 |
| Zc3h13 | -0.5126073 | 0.02410122 | 0.338535791 |
| Crlf2 | 0.692652924 | 0.02424246 | 0.339986375 |
| Atp7b | -1.003779129 | 0.02428165 | 0.340075309 |
| 1700025G04Rik | 2.23275423 | 0.02430801 | 0.340075309 |
| Slc44a1 | 0.617560666 | 0.02437165 | 0.340550848 |
| Exoc8 | -0.48055549 | 0.0244527 | 0.340604748 |
| Nrp2 | 2.688138575 | 0.02445935 | 0.340604748 |
| Ddx41 | -0.468179084 | 0.02448145 | 0.340604748 |
| Dnajc25 | -0.790555684 | 0.02450276 | 0.340604748 |
| Tuba1c | -0.48118901 | 0.02452378 | 0.340604748 |
| Mfap3l | 2.097179433 | 0.02458882 | 0.340610971 |
| Pglyrp2 | 3.333770589 | 0.02464285 | 0.340610971 |
| Mcc | 1.907008479 | 0.02476643 | 0.341907684 |
| Angpt1 | 1.259025115 | 0.02486301 | 0.342829028 |
| Samd11 | 1.269870089 | 0.02499911 | 0.344004794 |
| Cers4 | 1.174885292 | 0.02500818 | 0.344004794 |
| Hoxb3 | 1.928135988 | 0.02510713 | 0.344540549 |
| Mogs | -0.500287018 | 0.02536342 | 0.345870569 |
| Klk8 | 2.762674918 | 0.02539783 | 0.345870569 |
| Nab2 | 1.617574229 | 0.02548807 | 0.345870569 |
| Lpp | 1.836173808 | 0.02550246 | 0.345870569 |
| Il18r1 | 1.654277957 | 0.02550322 | 0.345870569 |
| Camk2d | 1.73731 | 0.02552989 | 0.345870569 |
| Sdk1 | -0.760649234 | 0.02555834 | 0.345870569 |

| | | | |
|---|---|---|---|
| Clcn2 | 0.897124664 | 0.02555912 | 0.345870569 |
| Faah | 1.440250558 | 0.02559613 | 0.345870569 |
| Gstk1 | 1.841414829 | 0.02563146 | 0.345870569 |
| Fam13a | 2.378863736 | 0.02566945 | 0.345870569 |
| Gnl1 | -0.469866032 | 0.02567407 | 0.345870569 |
| Rnaseh2a | -0.499886221 | 0.02571517 | 0.345870569 |
| Mcf2l | 1.438497679 | 0.02572446 | 0.345870569 |
| Ptpn21 | 0.787072989 | 0.02574607 | 0.345870569 |
| Tmem184b | 0.650352259 | 0.02588674 | 0.347354016 |
| Vezt | -0.714790417 | 0.02596265 | 0.34796618 |
| Ppard | 0.661788322 | 0.02602958 | 0.348456528 |
| Gja1 | -0.491779508 | 0.0261742 | 0.349270279 |
| Accs | 1.44897408 | 0.02626905 | 0.350030548 |
| Cfb | 1.384865865 | 0.02634997 | 0.350701927 |
| Grb10 | -0.44284699 | 0.02639813 | 0.350936267 |
| Acvr1b | 1.641913301 | 0.0265777 | 0.352507418 |
| A230046K03Rik | 0.476486026 | 0.02671168 | 0.353875899 |
| Serpind1 | 1.616769705 | 0.02677838 | 0.35405372 |
| Eid2 | 1.392034649 | 0.02678676 | 0.35405372 |
| Col8a1 | -0.452979927 | 0.02686478 | 0.354350119 |
| Kif2c | -0.529197791 | 0.02687964 | 0.354350119 |
| Plvap | -0.64114607 | 0.02693901 | 0.354350119 |
| Lrrc16a | 2.07505392 | 0.0269589 | 0.354350119 |
| Dlg2 | 1.90668922 | 0.02696343 | 0.354350119 |
| Aplf | -0.51180934 | 0.02701007 | 0.354557264 |
| Actn1 | 1.422588948 | 0.0270538 | 0.354725981 |
| Incenp | -0.631614312 | 0.02719379 | 0.355441979 |
| Cbarp | 0.635322049 | 0.02721311 | 0.355441979 |
| Osbpl6 | 1.710597223 | 0.02721883 | 0.355441979 |
| Tbccd1 | -0.527323985 | 0.02723219 | 0.355441979 |
| Nme1 | -0.464935216 | 0.02729033 | 0.35578283 |
| Mtss1l | 1.895456649 | 0.02732026 | 0.35578283 |
| Nedd9 | 1.200155491 | 0.02742506 | 0.35674318 |
| Lrrc66 | 1.591656283 | 0.02750379 | 0.35695881 |
| Lmnb1 | -0.55081411 | 0.02754133 | 0.357042559 |
| Hist1h4b | -0.648698519 | 0.02764882 | 0.357420821 |
| Zfp827 | 2.060751529 | 0.02790467 | 0.359009094 |
| Utp3 | -0.547596755 | 0.02791182 | 0.359009094 |
| Plac8 | 1.024005222 | 0.02800493 | 0.359803878 |
| Dnmt1 | -0.433136801 | 0.02804298 | 0.359890171 |
| E2f8 | -0.426659888 | 0.02821256 | 0.361274547 |
| Hipk2 | 0.608679467 | 0.02821376 | 0.361274547 |
| Plekha2 | 1.155657763 | 0.02839805 | 0.362579167 |

| | | | |
|---|---|---|---|
| Pik3r2 | -0.625961693 | 0.02841177 | 0.362579167 |
| St3gal4 | 0.474182947 | 0.02844244 | 0.362579167 |
| Plscr4 | 1.259066991 | 0.02848548 | 0.362579167 |
| Emc9 | -0.789771527 | 0.02851616 | 0.362579167 |
| Fli1 | 1.247841321 | 0.02852381 | 0.362579167 |
| Id3 | 2.715264379 | 0.02859625 | 0.362579167 |
| Naip2 | 0.585681809 | 0.02863526 | 0.362579167 |
| Ppp1r16b | 1.455747941 | 0.02866288 | 0.362579167 |
| Hist1h2ba | -0.929066061 | 0.02872817 | 0.36300521 |
| Fgl2 | 3.135527508 | 0.02902205 | 0.365522857 |
| Ncaph | -0.4719487 | 0.02908732 | 0.365522857 |
| Fam196b | 0.958031772 | 0.02910511 | 0.365522857 |
| Cdk18 | 1.97144111 | 0.02915051 | 0.365522857 |
| Calcrl | 0.909525292 | 0.02917186 | 0.365522857 |
| Tes | 1.515049819 | 0.0291882 | 0.365522857 |
| Vash1 | 2.081456806 | 0.02921649 | 0.365522857 |
| Rfxap | -0.533779984 | 0.02924565 | 0.365522857 |
| E2f4 | -0.494639152 | 0.0292991 | 0.365792922 |
| Pask | -0.508363193 | 0.02938865 | 0.366512477 |
| Epha4 | 3.089926817 | 0.02943717 | 0.366656386 |
| Bhlhe40 | 1.55358847 | 0.02946403 | 0.366656386 |
| Dmxl2 | 0.889993569 | 0.02952828 | 0.367058203 |
| Ccng2 | 1.07766353 | 0.02966884 | 0.368177948 |
| Cask | 0.77248908 | 0.02969939 | 0.368177948 |
| Adarb1 | 1.787158224 | 0.02972015 | 0.368177948 |
| Rtn2 | 1.699485581 | 0.02974657 | 0.368177948 |
| Grid1 | -0.640952458 | 0.02980658 | 0.368523579 |
| Trf | 3.04210992 | 0.0298956 | 0.368860734 |
| Gsn | 0.722269784 | 0.02989808 | 0.368860734 |
| Ankrd13d | 1.80984913 | 0.03005277 | 0.370213023 |
| Atp8b4 | 1.874287097 | 0.03007215 | 0.370213023 |
| Stard4 | 0.592396891 | 0.03021391 | 0.370832397 |
| Nek6 | 1.348880996 | 0.03024023 | 0.370832397 |
| Cep170b | 0.926741956 | 0.03025161 | 0.370832397 |
| Chd5 | 1.595922016 | 0.03031038 | 0.371156757 |
| Hirip3 | -0.690426721 | 0.03039231 | 0.371359316 |
| Cth | 1.708623691 | 0.03041448 | 0.371359316 |
| Smim3 | 1.08189326 | 0.03042392 | 0.371359316 |
| Ckap2l | -0.433525177 | 0.03048465 | 0.371705573 |
| Slc31a2 | 1.688624964 | 0.03063239 | 0.373028887 |
| Rnaseh2c | -0.517746101 | 0.03065813 | 0.373028887 |
| Trip13 | -0.601699649 | 0.03075692 | 0.373834872 |
| Bms1 | -0.557291976 | 0.03096467 | 0.3752113 |

| | | | |
|---|---|---|---|
| Pou4f1 | -0.469534132 | 0.03101542 | 0.3752113 |
| Ttll12 | -0.47741425 | 0.03102154 | 0.3752113 |
| Ptms | 1.056554601 | 0.03103356 | 0.3752113 |
| Hist1h1a | -0.837556084 | 0.03106616 | 0.3752113 |
| Fam101b | 1.645283656 | 0.03116253 | 0.375879189 |
| Ethe1 | 0.709877262 | 0.03120176 | 0.375879189 |
| Epor | -0.444543162 | 0.03156181 | 0.377723116 |
| Ckb | 1.452884883 | 0.0315652 | 0.377723116 |
| Fam214a | 0.829201702 | 0.0315811 | 0.377723116 |
| Extl3 | -0.424776594 | 0.03159087 | 0.377723116 |
| Pelp1 | -0.503583641 | 0.03164815 | 0.377723116 |
| Mrpl11 | -0.501953354 | 0.03168158 | 0.377723116 |
| Arl5b | 0.918730478 | 0.03170785 | 0.377723116 |
| Ufsp1 | 1.336209936 | 0.0317103 | 0.377723116 |
| Lzts3 | 1.512535374 | 0.0317312 | 0.377723116 |
| Sgol2a | -0.515744199 | 0.03173896 | 0.377723116 |
| Arhgap11a | -0.509159015 | 0.03176742 | 0.377723116 |
| Tmem184a | 1.931246393 | 0.03185623 | 0.378387458 |
| Slc35b3 | 0.680810764 | 0.03202843 | 0.379156256 |
| Ifitm1 | 1.897648705 | 0.03206961 | 0.379156256 |
| Gtf2h4 | -0.669971762 | 0.03211989 | 0.379156256 |
| Rgs9bp | -0.781025317 | 0.03214915 | 0.379156256 |
| Fndc3b | 1.036605469 | 0.0321982 | 0.379156256 |
| Nav1 | 1.57217085 | 0.03219869 | 0.379156256 |
| Ly75 | 2.28370269 | 0.03221805 | 0.379156256 |
| Gemin6 | -0.76770144 | 0.03240379 | 0.380951846 |
| Elmo3 | 0.994177581 | 0.03246228 | 0.381071181 |
| Oasl2 | 0.999554992 | 0.03250063 | 0.381071181 |
| Rgs2 | 1.816804603 | 0.03251347 | 0.381071181 |
| Hsdl2 | -0.563673318 | 0.03258338 | 0.381112685 |
| Pik3c2b | 0.81650705 | 0.03270174 | 0.381753775 |
| Timeless | -0.579883923 | 0.03270466 | 0.381753775 |
| 2410016O06Rik | -0.455911808 | 0.03282715 | 0.381916712 |
| Sart3 | -0.566352745 | 0.03287687 | 0.381916712 |
| Arf4 | 0.426810079 | 0.03288726 | 0.381916712 |
| Ralb | 1.37356492 | 0.03288838 | 0.381916712 |
| Tnfaip2 | -0.468536334 | 0.03292703 | 0.381916712 |
| Prkaca | -0.439087859 | 0.03295137 | 0.381916712 |
| Wrnip1 | -0.494514424 | 0.03307972 | 0.38288986 |
| Tubb5 | -0.412632089 | 0.03310201 | 0.38288986 |
| Ttyh2 | 1.17412684 | 0.03316577 | 0.382964565 |
| Grasp | 2.314232926 | 0.03317515 | 0.382964565 |
| Tbxas1 | 2.034498856 | 0.03336186 | 0.384733229 |

| | | | |
|---|---|---|---|
| Ppl | 2.162876523 | 0.03343408 | 0.385179385 |
| Olig1 | -0.734591644 | 0.03354453 | 0.386064643 |
| Itga6 | 0.777412833 | 0.03362216 | 0.386570654 |
| Mctp1 | 1.823960271 | 0.03378209 | 0.388021135 |
| Faap24 | -0.632454444 | 0.0340061 | 0.389814444 |
| Nbea | -0.58094252 | 0.0340789 | 0.390122037 |
| Fzd5 | 0.7379289 | 0.03410086 | 0.390122037 |
| P2rx4 | 0.610039627 | 0.03415986 | 0.390408145 |
| Tjap1 | -0.525402278 | 0.03432436 | 0.391128086 |
| Usp3 | 0.467994211 | 0.03434023 | 0.391128086 |
| Matk | 0.785872312 | 0.0343442 | 0.391128086 |
| Grtp1 | 1.341133495 | 0.03439854 | 0.391128086 |
| Slc17a4 | 2.352122179 | 0.03440143 | 0.391128086 |
| Rbp1 | 1.287102721 | 0.03442717 | 0.391128086 |
| Mpl | 3.028516673 | 0.0346127 | 0.392847349 |
| Dhx16 | -0.538685415 | 0.03466973 | 0.392881487 |
| Scin | 1.161549747 | 0.03478278 | 0.392992385 |
| Zfp532 | 3.37252269 | 0.03479656 | 0.392992385 |
| Loxl2 | 0.668475586 | 0.03497627 | 0.394622143 |
| Sf3a3 | -0.466812253 | 0.03505468 | 0.394622143 |
| Glipr1 | 1.270372554 | 0.03506464 | 0.394622143 |
| Tbc1d32 | 1.037188452 | 0.03511014 | 0.394622143 |
| Slco4a1 | 2.445757196 | 0.0351301 | 0.394622143 |
| Gstm1 | 1.203049532 | 0.03518824 | 0.394622143 |
| Hk2 | 0.733135229 | 0.03520194 | 0.394622143 |
| Gtse1 | -0.515008675 | 0.03521572 | 0.394622143 |
| Megf11 | 0.943444043 | 0.03529783 | 0.395156804 |
| Fbxo32 | 1.488160049 | 0.03551393 | 0.396802563 |
| Mfsd6 | 0.886625766 | 0.03580071 | 0.399430611 |
| Slc6a8 | 1.553582168 | 0.0358187 | 0.399430611 |
| Mfsd14b | -0.415618221 | 0.03592558 | 0.400233914 |
| Crebrf | 0.504933426 | 0.03602763 | 0.400809675 |
| Lpar6 | 1.503378083 | 0.03606433 | 0.400809675 |
| Prr36 | 2.109648169 | 0.03608194 | 0.400809675 |
| Hjurp | -0.491691327 | 0.03614383 | 0.401066018 |
| Zfp1 | -0.64316568 | 0.03620845 | 0.401066018 |
| Vps54 | 0.426704972 | 0.03620977 | 0.401066018 |
| Psen2 | -0.450682608 | 0.03626245 | 0.401128566 |
| Synj2 | 0.999073088 | 0.03628881 | 0.401128566 |
| 2310022B05Rik | -0.490529298 | 0.03632019 | 0.401128566 |
| Lrmp | 0.577867587 | 0.0364197 | 0.401410402 |
| Camsap3 | 0.960884032 | 0.03647989 | 0.401410402 |
| Pibf1 | 0.473060154 | 0.0364855 | 0.401410402 |

| | | | |
|---|---|---|---|
| Tmem176b | 1.887115274 | 0.03659108 | 0.402186751 |
| Khsrp | -0.405608562 | 0.03674979 | 0.403545003 |
| 1700022I11Rik | 1.455074676 | 0.03690379 | 0.404660148 |
| Pcdhgc5 | 2.013676251 | 0.0369218 | 0.404660148 |
| Anxa4 | 1.044137467 | 0.03706067 | 0.405523449 |
| Fen1 | -0.559652139 | 0.03719472 | 0.40563977 |
| Ssx2ip | -0.601992119 | 0.03720737 | 0.40563977 |
| St8sia6 | 1.142977408 | 0.03722308 | 0.40563977 |
| Hip1 | 4.351496093 | 0.03730075 | 0.406100909 |
| Llgl2 | 0.737851696 | 0.03741014 | 0.40690612 |
| Diablo | -0.449550215 | 0.03750777 | 0.407582115 |
| Alpk1 | 0.762686413 | 0.03756665 | 0.407836017 |
| D030056L22Rik | -0.595874389 | 0.03774579 | 0.408771256 |
| Pnck | -0.7733866 | 0.03774738 | 0.408771256 |
| Nceh1 | 0.815351732 | 0.03776562 | 0.408771256 |
| Tonsl | -0.525865706 | 0.03780941 | 0.408771256 |
| Mpnd | 1.087506128 | 0.03785158 | 0.408771256 |
| Mcm2 | -0.472587302 | 0.03790191 | 0.408771256 |
| Leprot | -0.493820087 | 0.03796233 | 0.409038756 |
| Deptor | 0.60451775 | 0.03806809 | 0.409410152 |
| Arhgap23 | -0.567100216 | 0.03816563 | 0.409908205 |
| Nfkb1 | 0.406835374 | 0.03822786 | 0.409908205 |
| Cyth1 | 0.632398294 | 0.03824029 | 0.409908205 |
| Zfp946 | 1.159409012 | 0.03828472 | 0.409908205 |
| Pear1 | 1.980239153 | 0.03840371 | 0.409908205 |
| Mars | -0.441490458 | 0.03843968 | 0.409908205 |
| Suco | 0.457644824 | 0.03851685 | 0.409908205 |
| Hdac5 | 1.015063845 | 0.03856945 | 0.409908205 |
| Ier3 | 0.6895007 | 0.03858572 | 0.409908205 |
| Tsc22d3 | 0.659915779 | 0.03859173 | 0.409908205 |
| Klc4 | 0.914813144 | 0.03860032 | 0.409908205 |
| Nfe2l3 | 1.107786587 | 0.03860061 | 0.409908205 |
| Rab44 | 1.200918142 | 0.03860836 | 0.409908205 |
| Slc35f5 | 0.941822332 | 0.03869871 | 0.409908205 |
| Ddx23 | -0.500226266 | 0.03874782 | 0.409908205 |
| Pdcd7 | -0.476167022 | 0.03875093 | 0.409908205 |
| 1700061G19Rik | 1.39799158 | 0.03875692 | 0.409908205 |
| Mex3d | -0.539034133 | 0.03879246 | 0.409908205 |
| Arhgap27 | 1.427508622 | 0.03884067 | 0.410040358 |
| Fut8 | 0.445686316 | 0.03909921 | 0.41239079 |
| 2010111I01Rik | 0.548868146 | 0.03917438 | 0.412804533 |
| Tcof1 | -0.399669648 | 0.03924129 | 0.413130598 |
| Rab32 | 1.29681601 | 0.03933311 | 0.413211435 |

| | | | |
|---|---|---|---|
| Fut4 | 1.799746759 | 0.0393569 | 0.413211435 |
| Wdr76 | -0.435650892 | 0.03944216 | 0.413728406 |
| Dnajb1 | 0.489195557 | 0.03949728 | 0.413928608 |
| Tuba1b | -0.475384469 | 0.03954275 | 0.414027324 |
| Rara | 0.782128118 | 0.0395824 | 0.414042578 |
| Rragb | 0.804586644 | 0.0396163 | 0.414042578 |
| Ermap | -0.466760277 | 0.03971417 | 0.414688182 |
| Fndc3a | 0.734308858 | 0.03982002 | 0.415325709 |
| Per1 | 0.477233205 | 0.03984755 | 0.415325709 |
| Abca3 | -0.525050097 | 0.03996372 | 0.415456434 |
| Lama5 | 1.225469978 | 0.03999553 | 0.415456434 |
| Pde5a | 0.93608332 | 0.04000477 | 0.415456434 |
| Srpk3 | 0.876680243 | 0.04010898 | 0.415827223 |
| Stk39 | 1.813504251 | 0.04018279 | 0.415827223 |
| Rras2 | 1.34797598 | 0.04021847 | 0.415827223 |
| Sat1 | 0.501296532 | 0.04033761 | 0.415827223 |
| Card19 | 0.885136968 | 0.04034404 | 0.415827223 |
| Slc22a18 | 1.564889688 | 0.04036376 | 0.415827223 |
| Cltb | 0.739781603 | 0.04054856 | 0.417022111 |
| Esco1 | 0.449350968 | 0.04055515 | 0.417022111 |
| Fignl1 | -0.475211609 | 0.04062191 | 0.417022111 |
| Cln3 | 0.826121636 | 0.0406421 | 0.417022111 |
| Txnrd2 | -0.659913186 | 0.04069438 | 0.417022111 |
| Ctla2b | 2.541144683 | 0.04072841 | 0.417022111 |
| Ifi27 | -0.727180185 | 0.04073645 | 0.417022111 |
| Hexdc | 0.779269911 | 0.04120093 | 0.421401476 |
| Ociad2 | 1.256476208 | 0.0413426 | 0.422119178 |
| Slc8b1 | 1.056849734 | 0.04136048 | 0.422119178 |
| Kcng2 | -0.508543812 | 0.04138135 | 0.422119178 |
| Gtf2f1 | -0.679850631 | 0.04142855 | 0.422225693 |
| Parp14 | 0.497944215 | 0.04151045 | 0.422607831 |
| Kctd12 | 1.298405612 | 0.04153963 | 0.422607831 |
| Rbm42 | -0.401084324 | 0.04163376 | 0.423190537 |
| Tspan14 | 0.946698545 | 0.04171524 | 0.42364388 |
| Tcf3 | -0.411820422 | 0.04176704 | 0.42365876 |
| Bub1b | -0.450729076 | 0.04179047 | 0.42365876 |
| Mbnl2 | 0.626651919 | 0.04192966 | 0.424015262 |
| Cpne3 | 1.289769764 | 0.04194347 | 0.424015262 |
| Pusl1 | -0.621877421 | 0.04195992 | 0.424015262 |
| Hdgf | -0.563134066 | 0.04198398 | 0.424015262 |
| Ndrg2 | 1.054745688 | 0.04201022 | 0.424015262 |
| Pwwp2b | 0.912759535 | 0.04209696 | 0.424175616 |
| Gpsm2 | -0.524721773 | 0.04215656 | 0.424175616 |

| | | | |
|---|---|---|---|
| Uhrf1bp1 | -0.404180798 | 0.0421686 | 0.424175616 |
| Timm10 | -0.518154905 | 0.04227443 | 0.424175616 |
| Cyp51 | 0.531860284 | 0.04228074 | 0.424175616 |
| Prg4 | 1.497734062 | 0.04230529 | 0.424175616 |
| Dhrs3 | 1.865564945 | 0.04232559 | 0.424175616 |
| Pla2g4c | 0.660556198 | 0.04234177 | 0.424175616 |
| Pomgnt1 | -0.484635952 | 0.04241908 | 0.424175616 |
| Kif18b | -0.47108873 | 0.04242196 | 0.424175616 |
| Ypel1 | 0.981968634 | 0.04246359 | 0.424175616 |
| Luzp1 | 0.715270729 | 0.04247453 | 0.424175616 |
| Pole | -0.376614237 | 0.04250619 | 0.424175616 |
| Rbmxl1 | -0.44992464 | 0.04260955 | 0.424837969 |
| Plek2 | 1.374336705 | 0.04297956 | 0.427414002 |
| Psmd5 | -0.394276799 | 0.04310896 | 0.427917713 |
| Gnpnat1 | 0.566138419 | 0.04311624 | 0.427917713 |
| Arhgap15 | 0.82682782 | 0.04314627 | 0.427917713 |
| Arhgap31 | 1.383807091 | 0.04321913 | 0.427917713 |
| Chaf1a | -0.510728708 | 0.04322398 | 0.427917713 |
| Siva1 | -0.453054854 | 0.04325374 | 0.427917713 |
| Ccdc6 | -0.387794285 | 0.04340448 | 0.428698457 |
| Srgap3 | 1.440634791 | 0.04340731 | 0.428698457 |
| Melk | -0.581514106 | 0.04361154 | 0.430345506 |
| Dstn | 0.518816895 | 0.0437424 | 0.431266227 |
| Wdr81 | -0.467780622 | 0.04383249 | 0.43178387 |
| Lnx1 | 1.065078753 | 0.04388792 | 0.431959377 |
| Hspa12b | 0.875207213 | 0.04394562 | 0.432157047 |
| Mrpl55 | -0.793134607 | 0.04405701 | 0.43221819 |
| Snx3 | -0.414532151 | 0.0441136 | 0.43221819 |
| Loxl2 | 0.630336271 | 0.04412485 | 0.43221819 |
| Dusp3 | 1.265725697 | 0.04420263 | 0.43221819 |
| Dpp9 | -0.391322519 | 0.04420477 | 0.43221819 |
| Gabpb2 | -0.363766765 | 0.04423018 | 0.43221819 |
| Suds3 | -0.426379183 | 0.04423312 | 0.43221819 |
| Slx4 | -0.50368689 | 0.04425288 | 0.43221819 |
| Isyna1 | -0.377182181 | 0.04432566 | 0.43256122 |
| Rab3d | 0.852980258 | 0.04444287 | 0.433210155 |
| Rrnad1 | 0.541762365 | 0.04448194 | 0.433210155 |
| Tacc2 | 0.936461441 | 0.04450531 | 0.433210155 |
| Casp1 | 1.601312228 | 0.04464504 | 0.43400438 |
| Tpx2 | -0.527658801 | 0.04470691 | 0.43400438 |
| Hus1 | -0.42768588 | 0.04471394 | 0.43400438 |
| Nudc | -0.612046097 | 0.04474666 | 0.43400438 |
| Man2a2 | 0.811234255 | 0.04479382 | 0.43400438 |

| | | | |
|---|---|---|---|
| Igsf6 | 1.252680586 | 0.04508076 | 0.435453804 |
| Ppp1r13b | 0.547154867 | 0.04509042 | 0.435453804 |
| Ccdc134 | 0.69737422 | 0.04514389 | 0.435453804 |
| Phf13 | 1.363828891 | 0.04515284 | 0.435453804 |
| BC030867 | -0.604377372 | 0.04531726 | 0.43667282 |
| Mfhas1 | -1.01839499 | 0.04555228 | 0.438569562 |
| Spty2d1 | 0.415228862 | 0.04563673 | 0.438896129 |
| Rab11fip1 | 0.381563534 | 0.04566262 | 0.438896129 |
| Rogdi | 1.68747985 | 0.04578175 | 0.439303245 |
| Psmf1 | -0.478108471 | 0.04579941 | 0.439303245 |
| Capn12 | 1.498050924 | 0.04581972 | 0.439303245 |
| Ercc6l | -0.427037451 | 0.04590585 | 0.439395537 |
| Lonrf1 | -0.472956962 | 0.04597629 | 0.4396775 |
| Lsmem1 | 1.399050716 | 0.046109 | 0.439873734 |
| Hook1 | 0.51971859 | 0.04615386 | 0.439936334 |
| Eps8l1 | 1.34214463 | 0.04642161 | 0.441755278 |
| Nfkbid | 0.89811937 | 0.04648737 | 0.441797173 |
| Srgn | 0.442483116 | 0.04651344 | 0.441797173 |
| Nefh | -0.559713697 | 0.04654517 | 0.441797173 |
| Meis1 | 1.50694601 | 0.04657987 | 0.441797173 |
| Sema3b | 1.294570623 | 0.04669687 | 0.442462649 |
| Galnt11 | 1.014507376 | 0.04681023 | 0.442462649 |
| Vasp | 0.47375468 | 0.04694281 | 0.442521283 |
| Appl1 | 0.627931008 | 0.0469507 | 0.442521283 |
| Pcid2 | -0.415794321 | 0.04704869 | 0.442521283 |
| Ica1 | 0.80722006 | 0.04706412 | 0.442521283 |
| Bcl2 | 1.003703146 | 0.04706644 | 0.442521283 |
| Sox12 | -0.417390171 | 0.04708001 | 0.442521283 |
| Maml3 | 0.601154969 | 0.04724595 | 0.443717919 |
| Arhgap29 | 1.38478494 | 0.0472891 | 0.443760245 |
| Fbxo5 | -0.473041835 | 0.0473361 | 0.443838726 |
| Satb1 | 1.541387315 | 0.04741704 | 0.443879964 |
| Hist1h3f | -0.610452581 | 0.047436 | 0.443879964 |
| B4galt3 | -0.492997188 | 0.04745643 | 0.443879964 |
| Litaf | 0.507529158 | 0.04764751 | 0.444630348 |
| 2810459M11Ril | -0.399819119 | 0.04768391 | 0.444630348 |
| D2Wsu81e | -0.412807071 | 0.0476926 | 0.444630348 |
| Dock6 | 1.082611896 | 0.04769329 | 0.444630348 |
| Fxyd5 | 1.483400657 | 0.04773588 | 0.444630348 |
| Lrwd1 | -0.375496557 | 0.0477856 | 0.444630348 |
| Vps4a | -0.427815106 | 0.04780763 | 0.444630348 |
| Kif13b | 0.40113778 | 0.04792459 | 0.445181281 |
| Stard7 | -0.413367903 | 0.04796671 | 0.445181281 |

| | | | |
|---|---|---|---|
| Nfat5 | 0.469453954 | 0.04804949 | 0.445181281 |
| Txnl4b | 0.932476323 | 0.04805957 | 0.445181281 |
| Ctdp1 | -0.430334606 | 0.0480796 | 0.445181281 |
| Casc5 | -0.396401949 | 0.04809942 | 0.445181281 |
| Bin1 | 1.12607762 | 0.04818613 | 0.445624702 |
| Rad9b | 0.881907651 | 0.04824539 | 0.445813787 |
| Epha1 | 1.00588494 | 0.04844563 | 0.447247354 |
| Vsig10 | 1.152122777 | 0.04847841 | 0.447247354 |
| Ddx59 | -0.707579209 | 0.04854769 | 0.447527119 |
| Casp7 | 0.511698999 | 0.04860065 | 0.447656054 |
| Rpia | -0.485695828 | 0.04871569 | 0.448313959 |
| Scamp4 | -0.460441306 | 0.04875014 | 0.448313959 |
| Osbpl3 | 0.812013742 | 0.04897575 | 0.449245393 |
| Zbtb46 | -0.4775617 | 0.04903245 | 0.449245393 |
| Impact | 0.508926535 | 0.04905157 | 0.449245393 |
| Rfc1 | -0.354100106 | 0.04915926 | 0.449245393 |
| Sptbn2 | 1.106582105 | 0.04916601 | 0.449245393 |
| Irf8 | 1.946394911 | 0.04918145 | 0.449245393 |
| Slc6a20b | 1.344325405 | 0.04920344 | 0.449245393 |
| Tubg1 | -0.424333622 | 0.04929263 | 0.449345343 |
| Mfsd13a | 0.946332451 | 0.04951107 | 0.450978722 |
| Cc2d2a | 1.640287494 | 0.0496647 | 0.452019644 |
| Pcbp1 | -0.461280186 | 0.05005621 | 0.454638095 |
| Pnp | -0.429450051 | 0.05006601 | 0.454638095 |
| Mrpl44 | -0.549339331 | 0.05008081 | 0.454638095 |
| Ache | 0.746853541 | 0.05011073 | 0.454638095 |
| Gnrh1 | 1.545508277 | 0.05021934 | 0.455028644 |
| Parp6 | 0.551526744 | 0.05023301 | 0.455028644 |
| Sulf2 | 1.218833915 | 0.05037622 | 0.455966303 |
| Frat2 | -0.380013699 | 0.05060651 | 0.457690023 |
| Naa40 | -0.35721763 | 0.05067415 | 0.457861055 |
| Nsrp1 | -0.429181545 | 0.0507136 | 0.457861055 |
| Adgrg1 | 0.660930703 | 0.05078426 | 0.457861055 |
| Ap1b1 | -0.383911324 | 0.05078487 | 0.457861055 |
| Pnp2 | -0.583456303 | 0.05095589 | 0.458318269 |
| Hspg2 | 0.959755178 | 0.05097801 | 0.458318269 |
| Smc1a | -0.425412716 | 0.05099358 | 0.458318269 |
